# The proteasome acts as a hub for local and systemic plant immunity in *Arabidopsis thaliana* and constitutes a virulence target of *Pseudomonas syringae* type-III effector proteins

**DOI:** 10.1101/053504

**Authors:** Suayib Üstün, Arsheed Sheikh, Selena Gimenez-Ibanez, Alexandra Jones, Vardis Ntoukakis, Frederik Börnke

**Affiliations:** Plant Metabolism Group, Leibniz-Institute of Vegetable and Ornamental Crops (IGZ), Theodor-Echtermeyer-Weg 1, 14979 Großbeeren; Institut of Biochemistry and Biology, University of Potsdam, Karl-Liebknecht-Str. 24-25, Haus 29, 14476 Potsdam; School of Life Sciences, University of Warwick, Coventry CV4 7AL, UK; Plant Molecular Genetics Department, Centro Nacional de BiotecnologÍa-CSIC (CNB-CSIC), 28049 Madrid, Spain.

## Abstract

Recent evidence suggests that the ubiquitin-proteasome system (UPS) is involved in several aspects of plant immunity and a range of plant pathogens subvert the UPS to enhance their virulence. Here, we show that proteasome activity is strongly induced during basal defense in Arabidopsis and mutant lines defective in proteasome subunits *RPT2a* and *RPN12a* support increased bacterial growth of virulent *Pseudomonas syringae* DC3000 (*Pst*), strains in local leaves. Both proteasome subunits are required for PTI events such as production of reactive oxygen species and mitogen-activated protein kinases signaling as well as for defense gene expression. Furthermore, analysis of bacterial growth after a secondary infection of systemic leaves revealed that the establishment of systemic-acquired resistance (SAR) is impaired in proteasome mutants, suggesting that the proteasome plays an important role in defense priming and SAR. In addition, we show that *Pst* inhibits proteasome activity in a type-III secretion dependent manner. A systematic screen for type-III effector proteins from *Pst* for their ability to interfere with proteasome activity revealed HopM1, HopAO1, HopA1 and HopG1 as candidates. Identification of proteins interacting with HopM1 by mass-spectrometry indicate that HopM1 resides in a complex together with several E3 ubiquitin ligases and proteasome subunits, supporting the hypothesis that HopM1 associates with the proteasome leading to its inhibition. We conclude that the proteasome is an essential component of the plant immune system and that some pathogens have developed a general strategy to overcome proteasome-mediated defense.

**One sentence summary:** The proteasome is required for local and systemic immune responses and is targeted by Pseudomonas type-III effectors

## Introduction

The ubiquitin-proteasome system (UPS) is one of the main protein degradation systems of eukaryotic cells that not only removes misfolded and defective proteins but also controls various cellular pathways through the selective elimination of short-lived regulatory proteins (Vierstra, 2009). The UPS regulates many fundamental cellular processes, such as protein quality control, DNA repair and signal transduction (Sadanandom et al., 2012). Selective protein degradation by the UPS proceeds from the ligation of one or more ubiquitin proteins to the ε-amino group of a lysine residue within specific target proteins catalysed by the consecutive action of E1, E2, and E3 enzymes. The resulting ubiquitinated proteins are then recognized and degraded by the 26S proteasome. The 26S proteasome itself is a 2.5 MDa ATP-dependent protease complex composed of 31 subunits divided into two types of subcomplexes, namely the 20S core protease (CP) and the 19S regulatory particles (RPs). While the CP is a broad spectrum ATP-and ubiquitin-independent protease complex, the RP subcomplex assists in recognizing ubiquitinated target proteins and in opening the channel of the CP to insert the unfolded substrates into the CP chamber for degradation (Smalle and Vierstra, 2004). During the past few years, several studies have revealed that the UPS controls various processes in almost all aspects of plant homeostasis, comprising cell division, plant development, responses to plant hormones as well as abiotic and biotic stress responses (Sadanandom et al., 2012).

It is becoming increasingly obvious that the regulated protein turnover via UPS controls multiple aspects of plant immunity, including pathogen recognition, immune receptor accumulation, and downstream defence signalling (Marino et al., 2012). Plant immunity relies on a multi-layered system to detect and resist attempted pathogen invasion. Cell surface immune receptors recognize conserved pathogen-associated molecular patterns (PAMPs) and initiate basal defences, known as PAMP-triggered immunity (PTI) (Jones and Dangl, 2006). This recognition results in the initiation of intracellular down-stream signalling that leads to the production of reactive oxygen species (ROS), activation of mitogen-activated protein kinase (MAPK) cascades, transcriptional reprogramming, expression of pathogen-related (PR) proteins and callose deposition at the cell wall (Boller and Felix, 2009). Adapted plant pathogens are able to overcome PTI by delivering effector proteins into host cells and induce effector-triggered susceptibility (ETS). On the other hand, resistant plants have evolved the ability to monitor the presence or activities of effectors by intracellular immune receptors, commonly referred to as "resistance (R) proteins", resulting in effector-triggered immunity (ETI) (Jones and Dangl, 2006). ETI is often accompanied by the hypersensitive response (HR), a form of localized programmed cell death (PCD) at the primary infection site (Hofius et al., 2007), thereby restricting the pathogen spread within infected tissue.

Localised pathogen attack can also lead to increased resistance towards secondary infection in uninfected parts of the plants. This type of increased resistance is referred to as systemic acquired resistance (SAR) (Fu and Dong, 2013). After SAR has been induced, plants are primed (i.e. sensitized) to respond more rapidly and more effectively to a secondary infection. Long distance signalling between the primary infected leaf and distal leaves is required for the onset of SAR. The defence hormone salicylic acid (SA) is known to be critical for the establishment of SAR in remote tissue as it is supposed to induce SAR-related gene expression via the downstream regulator NON-EXPRESSER OF PR GENES1 (NPR1), a transcriptional co-activator (Fu and Dong, 2013). However, other signalling metabolites such as pipecolic acid (Navarova et al., 2012) have also been shown to play an essential role in the establishment of SAR and pipecolic acid signalling appears to presuppose effective SA-signalling (Bernsdorff et al., 2016).

The intricate molecular processes underlying the cellular changes during PTI, ETI and SAR require a high degree of proteomic plasticity, likely involving the UPS at various levels. For instance, recent studies identified that members of the U-box E3 ligase family are negative regulators of PTI (Trujillo et al., 2008; Stegmann et al., 2012). In addition, some key plant defence signalling components are degraded by the 26S proteasome pathway, including the PAMP receptor FLS2 (Lu et al., 2011), the master regulator of SA-dependent defence NPR1 (Spoel et al., 2009), and WRKY45 from rice. The latter is a BTH-inducible transcription factor conferring strong resistance to fungal blast (Matsushita et al., 2012). Apart from its function in regulating the turnover of components implicated in plant immunity, several proteasome components have been identified to directly contribute to defence responses such as ROS production and HR formation (Marino et al., 2012). In particular, PBA1, the catalytic subunit of the 20S has been proposed to act as a caspase-like enzyme during the induction of programmed cell death in response to avirulent bacterial strains (Hatsugai et al., 2009). Concomitant with the role of 20S subunits in plant immunity, RPN1a, a component of the RP, has been described to be required for resistance in *Arabidopsis* against biotrophic fungi (Yao et al., 2012). The latter study also showed that accumulation of RPN1a is affected by SA and that the *rpn1a* mutant has defects in SA accumulation upon infection with Pst. However, based on the analysis of additional mutants it appears that not all proteasome subunits play a similar role in immunity (Yao et al., 2012). Considering the role of the UPS in plant defence responses, co-evolution between pathogens and their respective host plants has selected for virulence factors that can manipulate the UPS by targeting or exploiting certain UPS components to enhance virulence during plant-pathogen interactions (Dudler, 2013). Gram-negative bacterial pathogens use a type III secretion system to inject so called type III effector (T3E) proteins into host cells to interfere with host cellular functions and immunity (Macho, 2016). Several T3Es from different genera of plant pathogenic bacteria such as *Pseudomonas* or *Xanthomonas* were shown to suppress plant defences by acting as E3 ligases (e.g. AvrPtoB, XopL), or by promoting ubiquitination and degradation of target proteins (e.g. HopM1)(Nomura et al., 2006; Singer et al., 2013; Üstün and Börnke, 2014; Banfield, 2015; Üstün and Börnke, 2015). A more direct way to subvert the UPS is achieved by SylA, a secreted small non-ribosomal peptide from *P. syringae* pv. *syringae*, which binds to the catalytic subunits of the 26S proteasome to inhibit its activity and suppress plant immune reactions, including stomatal closure and salicylic acid (SA)-mediated signalling (Groll et al., 2008; Schellenberg et al., 2010; Misas-Villamil et al., 2013). The first bacterial T3Es that directly target the proteasome for defence suppression are XopJ from *Xanthomonas campestris* pv. *vesicatoria* and HopZ4 from *Pseudomonas syringae*. Both closely related T3Es interact with the proteasomal component RPT6, a subunit of the 19S RP, to inhibit proteasome activity (Üstün et al., 2013; Üstün et al., 2014; Üstün and Börnke, 2015). In effect, this results in impaired turnover of the SA master regulator NPR1 and the attenuation of SA-dependent defence responses.

Despite the significance advances in the recent years, we still lack knowledge of how the proteasome is involved in defence responses and whether targeting the host proteasome is a general strategy of various plant pathogens to establish disease. In order to gather more information on this aspect of plant-bacteria interactions, we systematically analysed the contribution of the proteasome to plant immunity, in particular during local and systemic defence responses of *Arabidopsis thaliana* towards virulent *Pseudomonas syringae* DC3000 (*Pst*) bacteria. Furthermore, the ability of *Pst* to modulate proteasome function through the delivery of T3Es was investigated. Our results how that proteasome subunits RPT2a and RPN12a are required for PTI events and the establishment of SAR, being essential for resistance against pathogenic and non-pathogenic bacteria. In turn, we could reveal by a systematic screen of the T3E repertoire of *Pst* that this pathogen evolved T3Es to interfere with proteasome function. In particular, we could identify T3Es HopM1, HopAO1, HopA1 and HopG1 as candidates suppressing proteasome activity. Further biochemical analysis revealed that HopM1 interacts with multiple proteins including UPS related proteins. Based on the data presented, we conclude that the proteasome is an essential component of the plant immune system and pathogens developed the ability to overcome proteasome-mediated defence as a general virulence strategy.

## Results

### The proteasome is required during PTI and suppressed in a T3E-dependent manner

In order to investigate whether proteasome activity is modulated during induced defence in local leaves of the model plant *Arabidopsis thaliana* infected with *Pst*, proteasome activity was assessed in leaves of wild type *Arabidopsis* plants inoculated either with a *Pst* wild type strain or a *PstΔhrcC* strain, which is not able to deliver T3Es into the host cell. The measurements revealed that proteasome activity is significantly induced in *Pst ΔhrcC* infected leaves when compared to the mock control (Fig. 1A), while infection with *Pst* wild type bacteria leads to a significant reduction in activity. This indicates that proteasome activity is induced during basal defence in response to infection with the non-pathogenic bacterium *Pst ΔhrcC* and that virulent *Pst* can suppress this induction in a T3E - dependent manner. A western blot analysis using an antibody directed against ubiquitin on extracts from *Pst* wild type and *Pst ΔhrcC* infected leaves revealed the accumulation of ubiquitinated proteins in both the cases (Fig. 1B, upper panel). However, when probed with an antibody directed against the proteasome core subunit PBA1, only *Pst* wild type infected leaves showed accumulation of unprocessed PBA1 (Figure 1B, lower panel) indicating disturbed proteasome maturation (Book et al., 2010). These findings suggest that proteasome activity is induced as part of the defence response to non-pathogenic bacteria and also constitutes a virulence target for *Pst* during the compatible infection of *Arabidopsis*.

**Figure 1.**
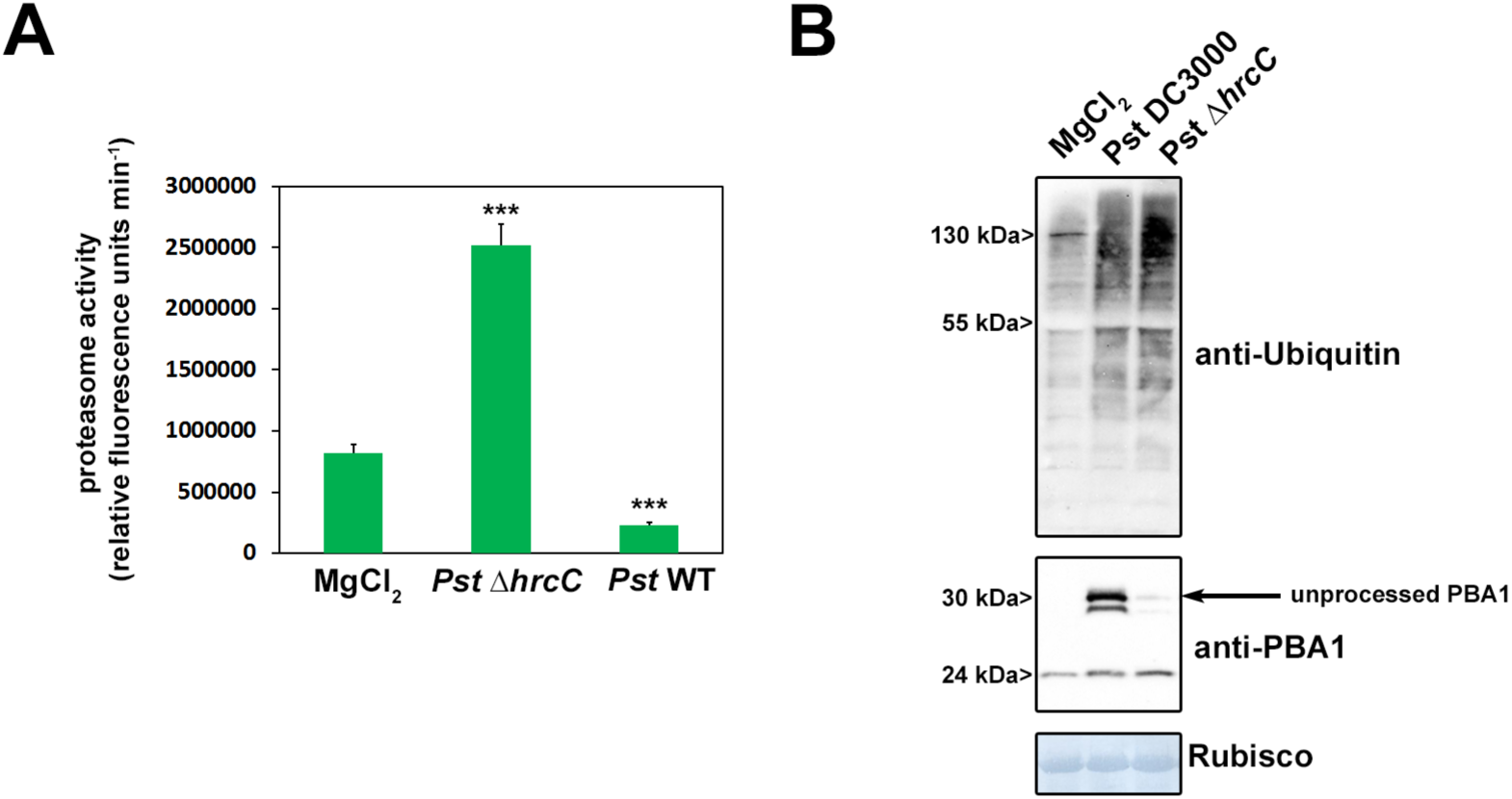
*Pseudomonas syringae* DC3000 prevents induction of proteasome activity during basal defense in a T3SS dependent manner. **(A)** Proteasome activity in leaves of *Arabidopsis* plants infected with either *Pst* wild type bacteria or infected with a *Pst* Δ*hrcC* strain lacking a functional T3SS. Samples were taken 2 dpi and the relative proteasome activity was determined. Each bar represents the mean of 3 biological replicates ± SD. MgCl_2_ infiltration serves as a mock control. Asterisks indicate a statistical difference according to student’s *t*-test (***, P < 0.001) in comparison to mock control. The experiment was repeated three times with similar results. **(B)** Accumulation of ubiquitinated proteins in *Arabidopsis* leaves after infection with different *Pst* strains (upper panel) and accumulation of the 20S subunit PBA1 (lower panel).

### Suppression of proteasome activity by *Pst* is independent of PTI inhibition and a functional SA signalling pathway

To rule out that the T3E-dependent suppression of early PTI during *Pst*-*Arabidopsis* interaction compromises proteasome function, we assessed whether the inhibition of the proteasome is linked to PTI suppression by T3Es. To this end, we infected plants with a *PstΔavrPto/avrPtoB* deletion strain that is compromised in suppression of early immune responses (He et al., 2006; Kvitko et al., 2009; Cunnac et al., 2011). In the absence of both T3Es *Pst* showed a similar degree of proteasome inhibition than the wild type (Fig. 2A), excluding the possibility that inhibition of early events of PTI is responsible for the effect on proteasome function and ruling out that neither AvrPto nor AvrPtoB are required for proteasome suppression.

**Figure 2.**
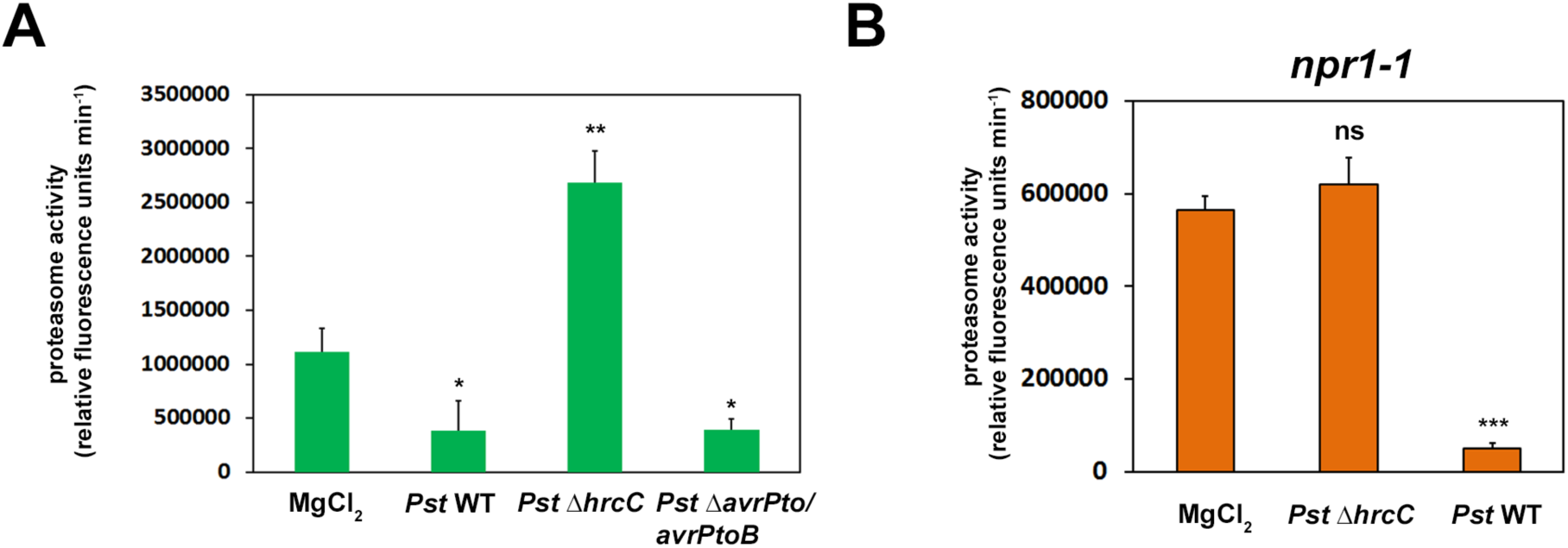
Suppression of proteasome induction by *Pseudomonas syringae* DC3000 is independent of the ability to suppress PTI and does not require a functional SA-signaling pathway. **(A)** Proteasome activity in *Arabidopsis* leaves infected with a *Pst* strain impaired in the inhibition of PTI *(Aavrpto/avrptoB)* compared to wild type and a T3SS deficient strain (*Pst* Δ*hrcC*). Samples were taken 2 dpi and the relative proteasome activity was determined. Each bar represents the mean of 3 biological replicates ± SD. MgCl_2_ infiltration serves as a mock control. Asterisks indicate a statistical difference according to student’s t-test (***, P < 0.001). **(B)** Proteasome activity in leaves of *npr1-1 Arabidopsis* plants 2 dpi with different *Pst* strains. Each bar represents the mean of 3 biological replicates ± SD. MgCl_2_ infiltration serves as a mock control. Asterisks indicate a statistical difference according to student’s t-test (***, P < 0.001).

Previous results demonstrated that the proteasome is activated by SA treatment and plants impaired in SA signalling are unable to induce proteasome activity upon infection (Gu et al., 2010; Üstün et al., 2013). In the light of these observations, we investigated whether the inhibitory effect of *Pst* on the proteasome is due to its ability to dampen SA signalling (DebRoy et al., 2004). To this end, we performed *Pst* infection assays with *npr1-1* mutant lines that lack a functional SA signalling (Cao et al., 1997). Consistent with previous experiments using wild type plants, *Pst ΔhrcC* was not able to induce proteasome activity in *npr1-1* (Fig. 2B), supporting the notion that proteasome activity seems to be at least partially induced by NPR1 - dependent SA-signalling during defence. However, *Pst* wild-type was still able to further inactivate the proteasome below levels detected in the mock control (Fig. 2B), demonstrating that *Pst* disables proteasome function independently of its ability to suppress SA signalling. This suggests that *Pst* directly inhibits proteasome activity, presumably by T3Es.

### Proteasome mutants are more susceptible to pathogenic and non-pathogenic *Pseudomonas* strains

In order to obtain genetic evidence for an involvement of the proteasome in basal defence responses a set of *Arabidopsis* mutant lines carrying defects in different proteasomal subunits was used for infection experiments with pathogenic bacteria. RPT2a is a subunit of the 19S RP of the proteasome where it gates the axial channel of the 20S core particle and controls substrate entry and product release. The *Arabidopsis rpt2a-2* mutant is a T-DNA insertion with only 25% of the RPT2 protein amount as compared to the wild type, which is due to the expression of the second RPT2 isoform encoded by the *RPT2b* gene (Lee et al., 2011). The *rpt2a-2* plants are affected in root elongation, leaf/organ size, trichome branching, endoreduplication, inflorescence stem fasciation, and flowering time (Lee et al., 2011). RPN12a is also part of the 19S RP where it is involved in complex assembly and the *Arabidopsis rpn12a-1* mutant was originally created by exon-trap mutagenesis and expresses an RPN12a-NPTII fusion protein whose incorporation into the 26S proteasome complex was proposed to have subtle effects on proteasome function (Smalle, 2002). The *rpn12a-1* mutant shows decreased rates of leaf formation, reduced root elongation, delayed skotomorphogenesis, and altered growth responses to exogenous cytokinins, suggesting that the mutant has decreased hormone sensitivity (Smalle et al., 2002). We first tested the susceptibility of both mutant genotypes towards virulent *Pseudomonas syringae* pv *maculicola* ES4326 (*Psm*) and monitored bacterial multiplication and symptom development 2 days post infection. As shown in Fig. 3A, *rpt2a-2* and *rpn12a-1* plants supported bacterial growth to significantly higher levels than the wild type and also showed accelerated symptom development on infected leaves (Fig. 3B). Thus, these data indicate that a fully functional proteasome is required to mount an efficient local defence response against virulent *Psm*. Infecting plants with the *Pst* strain also resulted in an enhanced bacterial proliferation in the proteasome mutants (Fig. 3C), which was also reflected by the stronger development of disease symptoms, such as leaf yellowing (Fig. 3D). Prompted by the finding that proteasome activity is highly induced upon *Pst*Δ*hrcC* infection, we next tested whether the proteasome mutant lines also show a higher sensitivity towards this strain. Measurement of bacterial growth revealed that the proteasome mutants supported more bacterial growth of the non-pathogenic *Pst*Δ*hrcC* (Fig. 3E). Thus the proteasome seems to play a critical role during PTI and is implicated in early immune responses.

**Figure 3.**
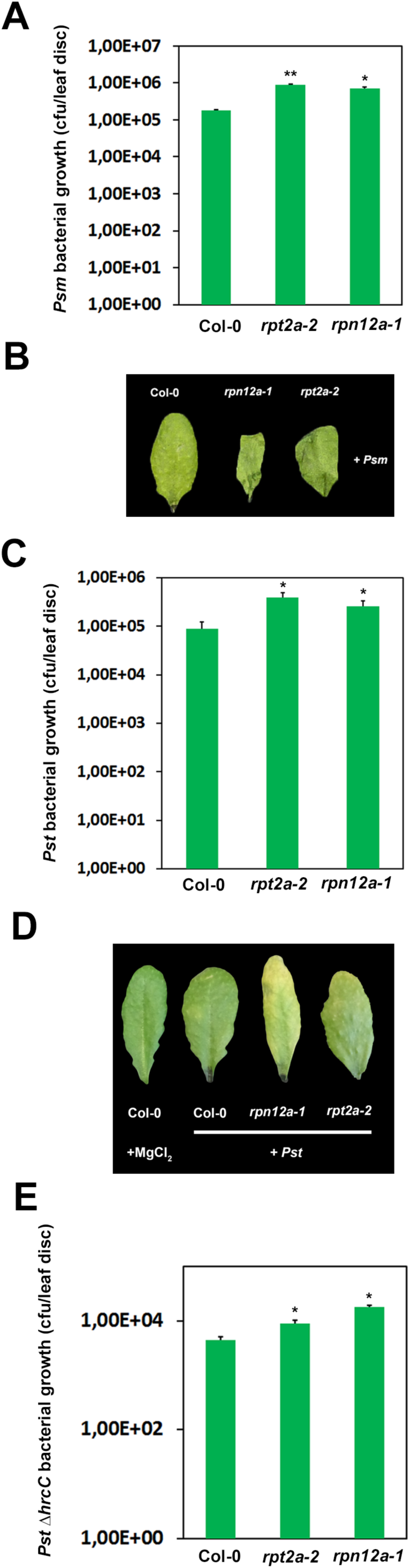
*Arabidopsis* mutant lines defective in different proteasome subunits display enhanced susceptibility towards infection with *Pst* DC3000 and *Psm*. **(A)** Bacterial density in leaves of different *Arabidopsis* genotypes infected with *Psm*. Leaves were syringe infiltrated with 1 x 10^5^ cfu/mL of bacteria and bacterial multiplication was determined at 2 dpi. Each bar represents the mean of 3 biological replicates ± SD. Asterisks indicate a statistical difference according to student’s t-test (**, P < 0.01; *, P < 0.05). **(B)** Phenotype of *Psm* infected *Arabidopsis* leaves 2 dpi. **(C)** Bacterial density in leaves of different *Arabidopsis* genotypes infected with *Pst* DC3000. Leaves were syringe infiltrated with a bacterial suspension of 1 x 10^5^ cfu/mL and *in planta* bacterial populations were determined 2 dpi. Each bar represents the mean of 3 biological replicates ± SD. Asterisks indicate a statistical difference according to student’s t-test (*, P < 0.05). **(D)** Phenotype of *Psm* infected *Arabidopsis* leaves 2 dpi. **(E)** Bacterial multiplication of an avirulent *Pst* Δ*hrcC* strain is enhanced in leaves of *Arabidopsis* proteasome mutant lines. Leaves were syringe infiltrated with a bacterial suspension of 1 x 10^5^ cfu/mL and bacterial multiplication was determined 2 dpi. Each bar represents the mean of 3 biological replicates ± SD. Asterisks indicate a statistical difference according to student’s t-test (*, P < 0.05).

### The proteasome is required for PAMP-triggered immunity

Because the proteasome is involved in local defence responses towards pathogenic and non-pathogenic bacteria, we further analysed the role of the proteasome during certain PTI responses. To this end, we first determined the production of reactive oxygen species. Recognition of flg22 by the PRR FLS2 (Gomez-Gomez and Boller, 2000) leads to an oxidative burst, one of the first measurable responses of plants to PAMP perception (Nicaise et al., 2009). ROS production in *rpt2a-2* and *rpn12a-1* plants was significantly attenuated compared to the Col-0 wild-type plants upon flg22 treatment (Fig.4A), indicating that early PTI responses such as ROS generation are partially dependent on a functional proteasomal turnover. In accordance with the decreased oxidative burst, the transcriptional response of *RbohD*, encoding an NADPH oxidase that is essential for flg22-triggered ROS production, was dampened in the proteasome mutant lines compared to Col-0 plants (Fig. 4B). Further downstream signalling cascades activated by the perception of PAMPs comprise the phosphorylation of MAPKs leading to the activation of MPK3, 6 and 4/11. Thus, we analysed whether this signalling cascade is altered in the proteasome mutants. Immunoblot analysis using an antibody against activated MAPKs revealed that in comparison to Col-0 plants, both proteasome mutants exhibited impaired kinetics of MAPK activation, as the phosphorylation signal was more rapidly fading out in the *rpt2a-2* and *rpn12a-1* mutants (Fig. 4C).

**Figure 4.**
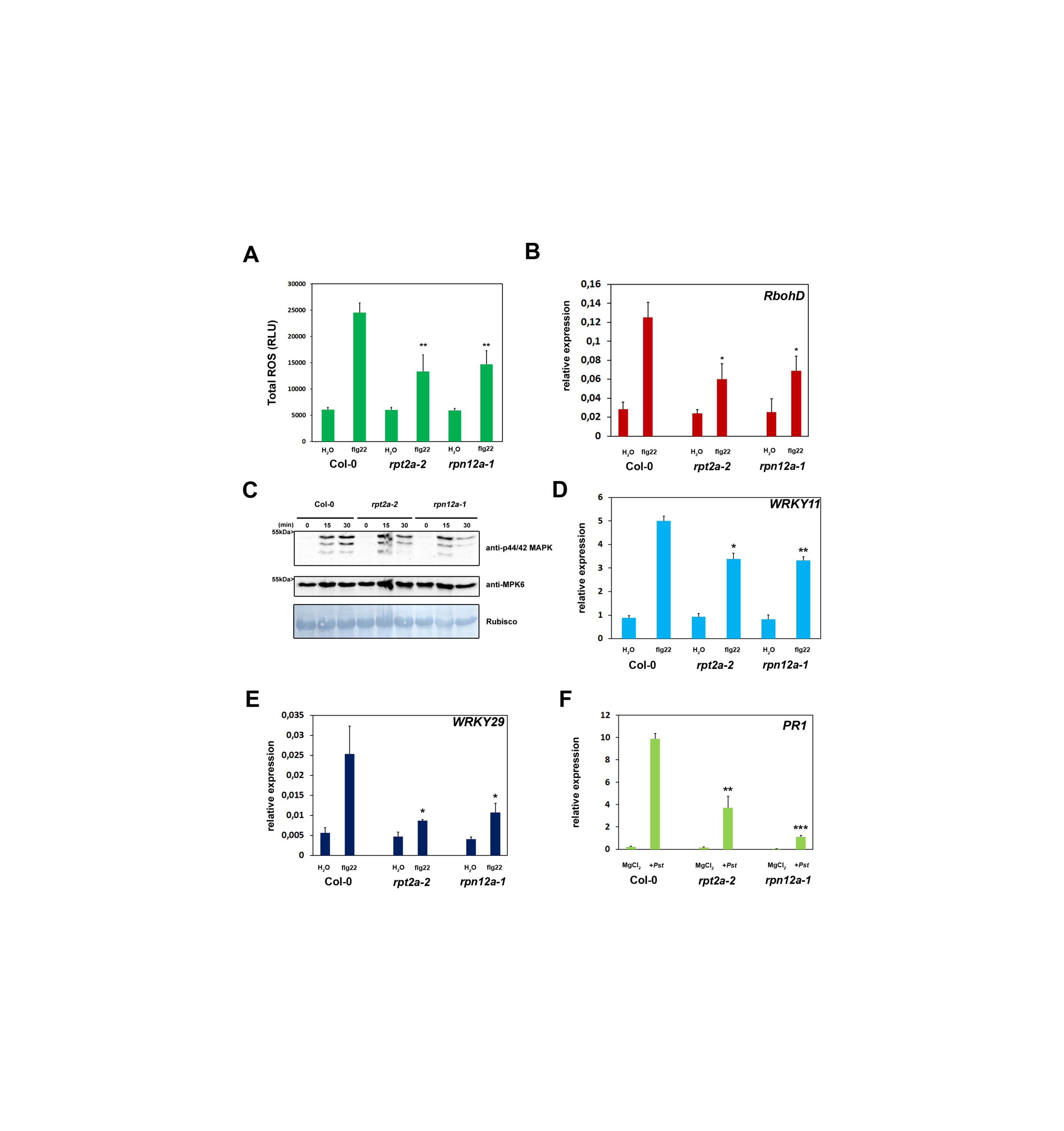
The proteasome is required for PAMP-Triggered Responses. **(A)** Total production of ROS in relative light units (RLU) during treatment with 1uM flg22 for 45 min. Each bar represents the mean of 4 biological replicates ± SD. Statistical significance compared with Col-0 plants treated with flg22 is indicated by asterisks (Student’s t test; **P < 0.01). ROS production was evaluated in at least two independent experiments with similar results. **(B)** Quantitative RT-PCR of ROS (*RboHD*) after flg22 treatment is reduced in proteasome mutants. Plants were treated with 1 μM flg22 or water (control). *UBC9* was used as a reference gene. Similar results were obtained in three independent experiments. Each bar represents the mean of 3 biological replicates ± SD. Changes in fold expression are significant for all genes in comparison to the wild type (+flg22) according to student’s t-test (**, P < 0.01; *, P < 0.05) **(C)** MAP Kinase signaling is comprised in proteasome mutant lines. Twelve-day-old seedlings were treated with 1 μM flg22 and samples were collected 0 to 30 min after treatment as indicated. Activated MAPKs were detected by immunoblotting using anti-p44/42 MAPK antibody. Proteins were also detected with anti-AtMPK6 antibody and amido black staining shows equal loading. Experiments were conducted twice with similar results. **(D, E)** PAMP-dependent induction of PTI marker genes is impaired in *Arabidopsis* proteasome mutant plants. Quantitative RT-PCR of immune response marker genes (*WRKY11* and *WRKY29*) 60 min after flg22 treatment. Plants were treated with 1 μM flg22 or water (control). *UBC9* was used as a reference gene. Similar results were obtained in three independent experiments. Each bar represents the mean of 3 biological replicates ± SD. Changes in fold expression are significant for all genes in comparison to the wild type (+flg22) according to student’s t-test (**, P < 0.01; *, P < 0.05). **(F)** Relative *PR1* expression is decreased in proteasome mutant lines in response to *Pst* infection. Plants were infiltrated with *Pst* and gene expression was analyzed 24hpi. Each bar represents the mean of 3 biological replicates ± SD. Changes in gene expression are significant for all genes in comparison to the wild type (+*Pst* infection) according to student’s t-test (**, P,0.01; ***, P,0.001).

In order to assess whether PTI is perturbed at the transcriptional level, we investigated the expression of PTI marker genes in *rpt2a-2* and *rpn12a-1* mutant plants and determined the gene expression of *WRKY11* and *WRKY29* upon flg22 stimulus (Stegmann et al., 2012). Real-time PCR revealed that the transcriptional activation of both PTI marker genes was significantly diminished in the proteasome mutant lines (Fig. 4D, E), providing further indications for a compromised PTI response in the proteasome mutants. Moreover, activation of *PR1* expression, a marker gene for the salicylic acid pathway, was also reduced in the proteasome mutants as compared to the control (Fig. 4F). Taken together, these data indicates that a fully functional proteasome is required during early and late PTI responses.

### Role of the proteasome in defence priming during SAR

Due to the fact that the expression of SA-inducible *PR1* was altered in the proteasome mutant lines (Fig.4F) and its prominent role during SAR, we investigated the role of the proteasome in defence priming during SAR. *Arabidopsis* proteasome mutants *rpt2a-2* and *rpn12a-1* were infected in lower (1°) leaves with the SAR-inducing pathogen *Psm* (Navarova et al., 2012). Analysis of bacterial growth after a secondary infection of systemic leaves (2°) two days after the primary infection revealed that bacterial multiplication was significantly inhibited in systemic leaves of wild type Col-0 plants, indicating that the primary infection with *Psm* induced effective SAR and primed the plants for a secondary infection of systemic leaves (Fig. 5A). In contrast, there was no significant difference in bacterial growth in systemic leaves of the proteasome mutants as compared to the unprimed mock control. Thus, the establishment of SAR is impaired in proteasome mutants as compared to Col-0 plants. This suggests that the proteasome plays an important role not only during local defence responses, but also in the establishment of systemic defence priming and SAR.

**Figure 5.**
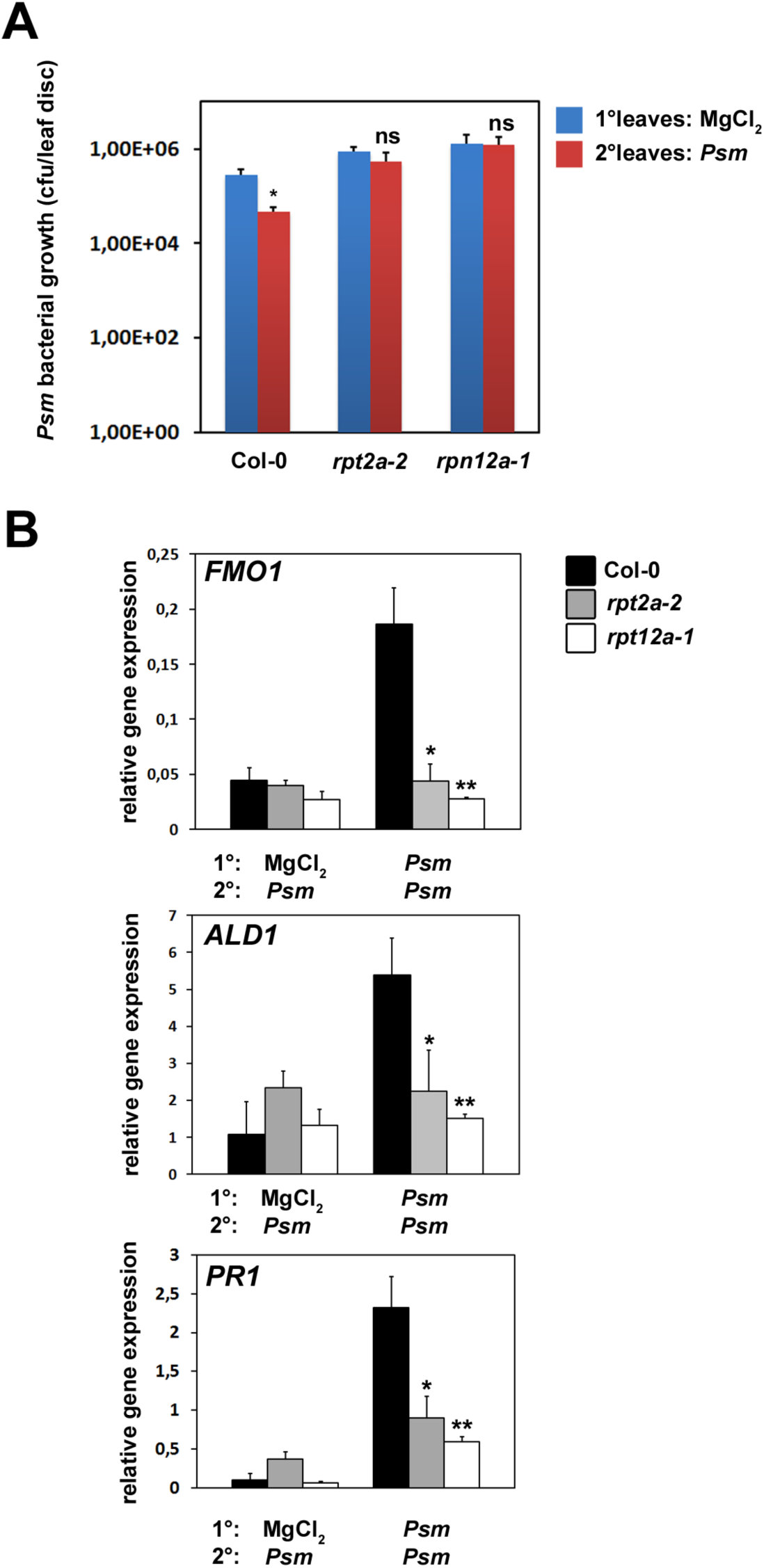
The proteasome is required for systemic acquired resistance. **(A)** SAR assay in Col-0, *rpt2a-2*, and *rpn12a-1* plants. Lower (1°) leaves were infiltrated with either 10 mM MgCl2 or *Psm* (OD 0.005), and 2 d later, three upper leaves (2°) were challenge infected with *Psm* (OD 0.001). Bacterial growth in upper leaves was assessed 2 d after 2° leaf inoculation. Bars represent the mean of 3 biological replicates ± SD. Asterisks denote statistically significant differences between indicated samples (* P < 0.05; ns, not significant; two-tailed t-test). **(B)** SAR priming of defense gene expression. Relative *ALD1, FMO1*, and *PR1* expression at 10 h after 2° treatment. Transcript levels were assessed by quantitative real-time PCR analysis, from three replicate samples (n=3). SD. Significant differences in comparions to Col-0 (*Psm*/*Psm* infected) were calculated using Student’s t-test and are indicated by: *, P,0.05; **, P,0.01.

To further corroborate this finding on the molecular level, SAR marker gene expression was analysed in systemic (2°) leaves of plants primed with *Psm* infection in local (1°) leaves. FLAVIN-DEPENDENT MONO-OXYGENASE1 (FMO1) is critical component of SAR establishment required for SA accumulation in systemic, noninoculated leaves (Mishina and Zeier, 2006) while AGD2-LIKE DEFENSE RESPONSEPROTEIN1 (ALD1) represents the aminotransferase required for the biosynthesis of the SAR signalling metabolite pipecolic acid (Navarova et al., 2012). As shown in Figure 5B, both *FMO1* and *ALD1* mRNA levels were strongly induced in 2° leaves of wild type *Arabidopsis* plants primed with *Psm* infection in 1° leaves as compared to the unprimed control, indicating that these plants effectively established SAR. On the contrary, *rpt2a-2* and *rpn12a-1* plants displayed no induction of *FMO1* or *ALD1* expression in challenge infected 2° leaves irrespective of the type of treatment of the 1° leaf. Thus, *rpt2a-2* and *rpn12a-1* plants lack induction of the essential SAR components FMO1 and ALD1 in systemic tissue upon a priming infection with *Psm* in 1° leaves. In order to analyses whether gene expression changes downstream of FMO1 and ALD1 are also impaired in *rpt2a-2* and *rpn12a-1* mutant plants, we determined the expression of the SA-responsive pathogenesis-related PR1 gene upon systemic infection of primed plants. The measurement revealed that *PR1* expression in systemic leaves of primed *rpt2a-2* and *rpn12a-1* plants was significantly reduced as compared to the wild type and displayed no induction in primed proteasome mutants as compared to the unprimed control (Figure 5B). Taken together the data indicate that a disturbance of proteasome function has repercussions on SAR-gene induction.

### *Pseudomonas* T3Es inhibit the proteasome

Given the finding that *Pst* suppresses proteasome activity in dependence of the delivery of T3Es, we further elucidated this phenomenon by setting up a fast screening system using a collection of *Pst* T3Es cloned into a binary plant expression vector for transient expression using Agrobacterium - infiltration. In total, we tested 16 *Pst* T3Es after transient expression in *Nicotiana benthamiana* for their ability to inhibit proteasome activity (Fig. 6A). Effector proteins AvrPto and AvrPtoB were excluded from this analysis, as previous assays using a *PstΔavrPto/avrPtoB* deletion strain showed that this strain was still able to compromise proteasome function (Fig. 2A). This experimental approach identified four T3Es (HopM1, HopG1, HopAO1 and HopA1) whose expression reproducibly led to almost 30 to 80% reduction in proteasome activity in *N. benthamiana* leaves (Fig. 6B). Notably, the inhibitory effect on the proteasome of HopM1, HopG1 and HopA1 in transient assays was even stronger than that of XopJ, a T3E from *Xanthomonas*, which was shown to inhibit the proteasome by degrading the RP subunit RPT6 (Üstün and Börnke, 2015). Expression of all T3Es tested was verified by western blot analysis using an anti-HA antibody (Fig. 6C). The remaining 12 T3Es from *Pst* tested were not able to suppress proteasome activity (Supp. Fig. S1-6), demonstrating that the interference with proteasome activity is specific for certain T3Es. This suggests that the proteasome might represent a virulence target for *Pst* and T3Es from *Pst* either directly or indirectly impede proteasome function.

**Figure 6.**
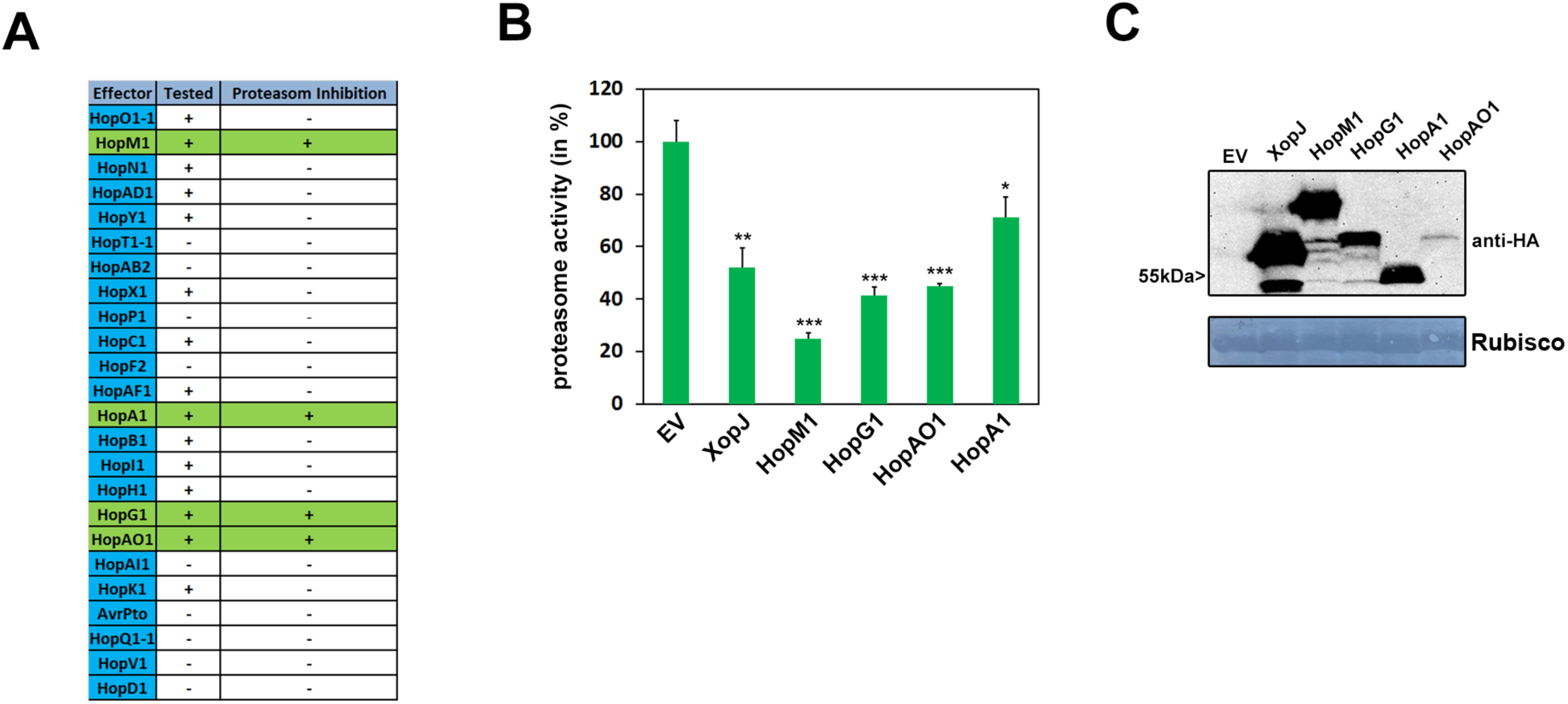
Type-III effectors from *Pst* DC3000 suppress proteasome activity in *N. benthamiana*. **(A)** Table of candidate effectors tested for their ability to modulate proteasome function. **(B)** Transient expression of T3Es HopM1, HopG1, HopAO1 and HopA1 in *Nicotiana benthamiana* inhibits proteasome activity. Proteasome activity in *N. benthamiana* leaves following transient expression of T3Es XopJ, HopM1, HopG1, HopA1, HopAO1 or empty vector control (EV). Relative proteasome activity in total protein extracts was determined by monitoring the breakdown of the fluorogenic peptide suc-LLVY-AMC at 30°C in a fluorescence spectrophotometer. EV was set to 100%. Data represent the mean ± standard deviation (SD) (n = 3). Asterisks indicate statistical significance (*P < 0.05, **P<0.01, ***P<0.001) determined by Student’s test (compared with EV control). **(C)** Protein extracts from *N. benthamiana* leaves transiently expressing T3Es tagged with HA and empty vector (EV) at 48 hpi were prepared. Equal volumes representing approximately equal protein amounts of each extract were immunoblotted and proteins were detected using anti-HA antiserum. Amido black staining served as a loading control.

### HopM1 inhibits proteasome activity during *Pseudomonas*-*Arabidopsis* interaction

Because HopM1 inhibits proteasome activity up to 80% and due to previously published work showing that HopM1 promotes degradation of its target protein AtMIN7 via the 26S proteasome (Nomura et al., 2006), we further concentrated our efforts on this candidate T3E. To demonstrate that HopM1 interferes with proteasome function during *Pst* infection of *A. thaliana*, we first infected plants with a *Pst* Δ*CEL* mutant strain that lacks 6 ORFs present in the Conserved Effector Locus (CEL), including the core T3Es AvrE and HopM1 (Alfano et al., 2000). The *Pst* Δ*CEL* mutant multiplies 200 to 500 fold less than wildtype *Pst* and fails to produce disease symptoms in Col-0 leaves. Complementation analysis suggests that ectopic HopM1 expression almost fully complements is mainly responsible for the *Pst* Δ*CEL* mutant phenotype (Nomura et al., 2006). Measurement of the overall leaf proteasome activity two days after infection reveals that leaves inoculated with *Pst* Δ*CEL* display proteasome activity that is comparable to mock control treated leaves (Fig. 7A), indicating that the lack of HopM1 renders *Pst* unable to inhibit proteasome activity during infection below the basal level but is still able to prevent its induction. Consistent with the activity data on the proteasome, accumulation of ubiquitinated proteins was also less pronounced compared to *Pst* wild-type infected leaves (Fig. 7B). Because this conserved effector locus also harbours AvrE, HopAA1-1, and HopN1 besides HopM1, (Kvitko et al., 2009) we decided to test a *Pst* strain that carries a deletion in HopM1 only, to exclude possible additional effects of the other T3Es. Infection of *Arabidopsis* plants with this strain provides evidence that *Pst* Δ*hopM1* is not able to reduce proteasome activity compared to the wild-type Pseudomonas strain (Fig. 7C). This is also partially reflected by a reduced accumulation of ubiquitinated proteins compared to *Pst* wild-infected plants (Fig. 7D). Thus, from this we could conclude that HopM1 is responsible for the inhibition of proteasome activity during the compatible interaction of *Pst* and *Arabidopsis*.

**Figure 7.**
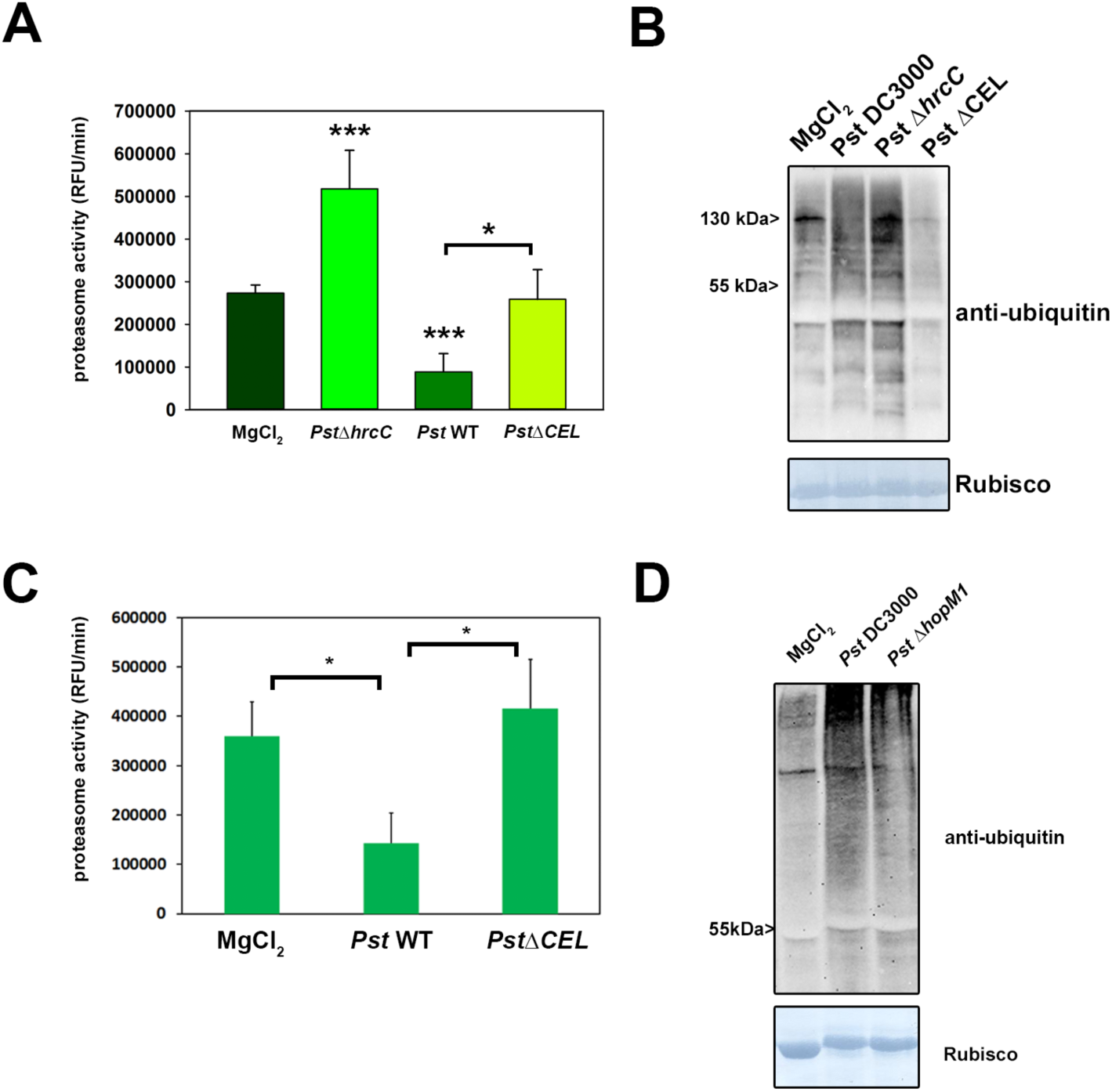
*Pseudomonas* T3E HopM1 is required for proteasome inhibition during *Pst* DC3000-*Arabidopsis* interaction. **(A)** Proteasome activity in leaves of *Arabidopsis* plants infected with either *Pst* wild type bacteria, *Pst ΔhrcC* and *Pst ΔCEL* strain lacking the conserved effector locus harboring HopM1. Samples were taken 2 dpi and the relative proteasome activity was determined. Each bar represents the mean of 3 biological replicates ± SD. MgCl_2_ infiltration serves as a mock control. Asterisks indicate a statistical difference according to student’s *t*-test (* P < 0.05; ***, P < 0.001). The experiment was repeated three times with similar results. **(B)** Accumulation of ubiquitinated proteins in *Arabidopsis* leaves after infection with different *Pst* strains indicated in the figure was determined using an anti-ubiquitin antibody. **(C)** Proteasome activity in leaves of *Arabidopsis* plants infected with either *Pst* wild type bacteria, *Pst* Δ*hrcC* and *Pst hopM1*. Samples were taken 2 dpi and the relative proteasome activity was determined. Each bar represents the mean of 3 biological replicates ± SD. MgCl_2_ infiltration serves as a mock control. Asterisks indicate a statistical difference according to student’s *t*-test (* P < 0.05). The experiment was repeated two times with similar results. **(D)** Accumulation of ubiquitinated proteins in *Arabidopsis* leaves after infection with different *Pst* strains indicated in the figure was determined using an anti-ubiquitin antibody.

### HopM1 interacts with components of the UPS *in-vivo*

To demonstrate the possible mechanism by which HopM1 recuces the total proteasome activity of the plant cell, an unbiased proteomics based screening was performed to find the *in vivo* interactions of HopM1 in *Nicotiana benthamiana*. HopM1 was immunoprecipitated and interacting proteins were identified using mass spectrometry (LC-MS/MS) analysis. The expression of HopM1 in *Nicotiana* was confirmed by anti-GFP western blot (Supp. Fig.S7). It was observed that many proteins related to the ubiquitin-proteasome system were associated with HopM1. The *Nicotiana* orthologs of known HopM1 interactors such as *Arabidopsis* AtMIN7 (a host ADP ribosylation guanine nucleotide exchange factor involved in membrane traffic), and AtMIN10 (a 14-3-3 protein) were also detected reflecting the interactions were specific to HopM1 (Nomura et al., 2006). After performing Fisher Exact test (p<= 0.05), a number of proteins related to 26S proteasome non-ATPase regulatory subunits 2, 3, 6, 12 and 14 were significantly enriched in the two independent screens carried out **(Table 1)**. E3 ubiquitin protein ligases UPL 1 and 3 were also significantly enriched in the screen further suggesting the role of HopM1 in directly perturbing the proteasome activity of the plant cell. However, the most interesting protein detected was proteasome-associated protein ECM29, which is known to stop the protein degradation by inhibiting the proteasomal ATPase activity in yeast (De La Mota-Peynado et al., 2013). The known target of direct ubiquitination post elicitation includes the flg22 receptor FLS2 (Lu et al., 2011). To analyse the possible effect of HopM1 induced decrease in proteasome activity on PTI signaling, the protein levels of FLS2 were analysed in *Arabidopsis* protoplasts transfected with HopM1. It was observed that the FLS2 levels accumulated in HopM1 expressing protoplasts without any PAMP elicitation in comparison to control. Significatly higher levels of FLS2 were also observed after longer (60 min) treatments with flg22 **(Fig. 8)**.

**TABLE 1:** List of UPS related proteins co-immunoprecipitated with HopML The significant values (Fisher Exact test p<=0.05) are indicated by Green color.

| Table 1: List of UPS related proteins co-immunoprecipitated with HopM1 |  |  |  | Total Unique Peptide Count |  |  |  |  |
| --- | --- | --- | --- | --- | --- | --- | --- | --- |
|  |  |  |  | GFP Control |  | HopM1 |  |  |
| Name of the Protein | Accession Number | Molecular Weight | Fisher's Exact Test (p <= 0.05) | Rep 1 | Rep 2 | Rep 1 | Rep 2 | At homologue |
| 26S Proteasome related proteins |  |  |  |  |  |  |  |  |
| Cluster of Proteasome associated protein ECM29 | NbS00044170g0008.1 SGN | 198 kDa | < 0.00010 | 1 | 0 | 18 | 22 | AT2G26780 |
| Cluster of 26S proteasome regulatory subunit | NbS00007460g0008.1 SGN [2] | 142 kDa | 0.023 | 8 | 1 | 14 | 14 | AT2G32730 |
| Cluster of Probable 26S proteasome non-ATPase regulatory subunit 3 | NICBE_198688.1 TGAC [3] | 55 kDa | < 0.00010 | 4 | 0 | 15 | 22 | AT1G75990 |
| Cluster of 26S protease regulatory subunit 6B homolog | NICBE_268659.1 TGAC [4] | 64 kDa | 0.08 | 6 | 0 | 8 | 7 | AT5G58290 |
| Cluster of 26S proteasome non-ATPase regulatory subunit 2 1A | NICBE_166405.1 TGAC [2] | 98 kDa | < 0.00010 | 6 | 0 | 19 | 21 | AT2G20580 |
| 26S protease regulatory subunit 7 homolog A | NICBE_154102.1 TGAC (+2) | 48 kDa | 0.0013 | 2 | 0 | 13 | 11 | AT1G53750 |
| Cluster of 26S protease regulatory subunit 6A homolog | NICBE_248120.1 TGAC [4] | 53 kDa | 0.018 | 2 | 0 | 12 | 3 | AT3G05530 |
| Cluster of 26S proteasome non ATPase regulatory subunit 11 | NbS00016436g0001.1 SGN [4] | 47 kDa | 0.08 | 3 | 0 | 9 | 4 | AT1G29150 |
| Cluster of 26S proteasome non-ATPase regulatory subunit 2 1A | NICBE_172900.1 TGAC [2] | 98 kDa | 0.0011 | 6 | 0 | 15 | 18 | AT2G20580 |
| Cluster of 26S proteasome non-ATPase regulatory subunit 12 | NICBE_244180.1 TGAC [2] | 51 kDa | 0.0069 | 1 | 0 | 10 | 7 | AT5G09900 |
| Probable 26S proteasome non-ATPase regulatory subunit 6 | NICBE_167161.1 TGAC (+1) | 44 kDa | 0.033 | 2 | 0 | 7 | 8 | AT4G24820 |
| 26S protease regulatory subunit 8 homolog A | NICBE_417606.1 TGAC (+1) | 47 kDa | 0.077 | 2 | 1 | 8 | 7 | AT5G19990 |
| Cluster of 26S protease regulatory subunit S10B homolog B | NICBE_194312.1 TGAC | 44 kDa | 0.13 | 3 | 0 | 7 | 6 | AT1G45000 |
| 26S proteasome non-ATPase regulatory subunit 14 | NICBE_364948.1 TGAC (+1) | 45 kDa | 0.039 | 0 | 0 | 6 | 2 | AT5G23540 |
| 26S proteasome non ATPase regulatory subunit 7 | NbS00041405g0012.1 SGN (+1) | 35 kDa | 0.48 | 3 | 0 | 2 | 6 | AT5G05780 |
| Ubiquitin related proteins |  |  |  |  |  |  |  |  |
| Cluster of Ubiquitin protein ligase 1 | NbS00002172g0012.1 SGN [6] | 319 kDa | < 0.00010 | 2 | 0 | 59 | 23 | AT1G55860 |
| Cluster of Auxin transport protein BIG | NbS00009154g0017.1 SGN [2] | 537 kDa | < 0.00010 | 4 | 0 | 41 | 16 | AT3G02260 |
| Cluster of E3 ubiquitin protein ligase UPL3 | NbS00031527g0002.1 SGN [3] | 199 kDa | 0.0048 | 1 | 0 | 16 | 1 | AT4G38600 |
| Cluster of E3 ubiquitin-protein ligase UPL1 | NICBE_298798.1 TGAC [2] | 389 kDa | 0.0016 | 0 | 0 | 7 | 9 | AT1G55860 |
| Ubiquitin conjugation factor E4 | NbS00018754g0010.1 SGN | 91 kDa | 0.26 | 3 | 0 | 8 | 3 | AT5G15400 |
| Cluster of Ubiquitin-conjugating enzyme E2 36 | NICBE_251503.1 TGAC [2] | 17 kDa | 0.11 | 1 | 0 | 8 | 0 | AT1G16890 |
| Cluster of E3 ubiquitin protein ligase listerin | NbS00000796g0017.1 SGN | 135 kDa | 0.039 | 0 | 0 | 2 | 6 | AT5G58410 |
| Probable ubiquitin conjugation factor E4 | NICBE_244276.1 TGAC | 83 kDa | 0.56 | 3 | 0 | 5 | 2 | AT5G15400 |
| Known HopM1 Interactors |  |  |  |  |  |  |  |  |
| Cluster of ARF guanine nucleotide exchange factor 2 | NbS00006288g0001.1 SGN [3] | 207 kDa | < 0.00010 | 5 | 0 | 28 | 29 | AT3G43300 (MIN7 like) |
| Cluster of Sec7 guanine nucleotide exchange factor | NbS00008405g0008.1 SGN [2] | 171 kDa | < 0.00010 | 5 | 0 | 35 | 46 | AT3G43300 (MIN7 like) |
| Sec7 guanine nucleotide exchange factor | NbS00016714g0003.1 SGN | 197 kDa | 0.0052 | 0 | 0 | 9 | 4 | AT3G43300 (MIN7 like) |
| Cluster of 14-3-3-like protein 16R | NICBE_421663.1 TGAC [4] | 29 kDa | 0.14 | 7 | 0 | 17 | 7 | AT4G24150 (MIN10 like) |

**Figure 8.**
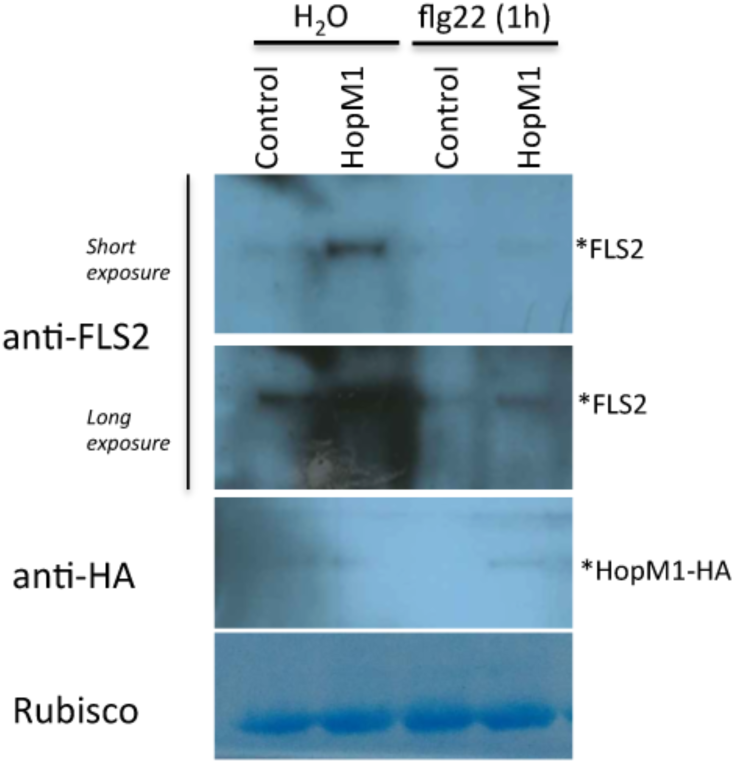
HopM1 leads to the accumulation of FLS2 protein. Western blot showing the protein levels in *Arabidopsis* Col-0 protoplasts transformed with either control or HopM1-HA using FLS2-specific antibody. The protoplasts were harvested after treating with water as control or 100 nM flg22 for 1 h.

## Discussion

Plant immunity has to be tightly regulated to ensure effective immune response activation with minimal negative effects on plant growth and reproduction. One such way to regulate certain immune processes is the proteasome-mediated recycling of defence components, highlighting an important role of the plant proteasome during plant immunity. Thus, evidence has accumulated that the 26S proteasome regulates plant defence responses at several layers of the surveillance system and hence constitutes a strategic target for plant pathogens to suppress host immune responses (Marino et al., 2012; Dudler, 2013).

In this study, we show that fully functional proteasome subunits RPT2a and RPN12a are required for the proper execution of PTI events in locally infected leaves and the establishment of SAR and thus, are essential for full resistance against *Pseudomonas* bacteria and also for the growth restriction of virulent bacteria in local and systemic tissue. Furthermore, we could show that *Pst* is able to inhibit proteasome activity during infection and that this phenomenon is dependent on T3E injection. A systematic screen of a *Pst* T3E collection identified candidate effector proteins for proteasome interference during a compatible interaction.

In the past, the proteasome has mainly been associated with the regulation of plant growth and development because the UPS plays an essential role in almost all aspects of hormone perception and signalling and many *Arabidopsis*proteasome subunit mutants show a range of developmental defects (Kurepa and Smalle, 2008). This is also true for the *Arabidopsis rpt2a-2* and *rpn12a-1* mutants used in the present study, both of them being part of the 19S RP of the proteasome. The *rpt2a-2* mutant is affected in root elongation, leaf/organ size, trichome branching, endoreduplication, inflorescence stem fasciation, and flowering time (Lee et al., 2011), while the *rpn12a-1* mutant shows decreased rate of leaf formation, reduced root elongation, delayed skotomorphogenesis, and altered growth responses to exogenous cytokinins (Smalle et al., 2002). However, it is important to mention that both mutants are characterised by weak defects on overall proteasome function since the RPT2 and RPN12 proteins are encoded by two almost identical genes and the second gene, albeit lower expressed, remains intact in the mutant plants (Kurepa and Smalle, 2008). Beyond their role in the regulation of plant growth and development certain proteasome subunits have been implicated to also play a role during plant immunity. For instance, a mutation in the 19S RP subunit RPN1a was identified as a suppressor of increased resistance to the adapted biotrophic powdery mildew pathogen *Golovinomyces cichoracearum* in *EDR2*(*Enhanced Disease Resistance 2*) loss of function mutants (Yao et al., 2012). The *Arabidopsis rpn1a* single mutant was more susceptible towards local infection with virulent and avirulent *P. syringae*strains as well as to *G. cichoracearum* (Yao et al., 2012). Infected *rpn1a* plants displayed a reduction in late defence responses such as the accumulation of the defence hormone salicylic-acid (SA) and a reduced expression of the defence marker gene *PR1*. The observation that the *rpt2a* as well as the *rpn12a* mutant show a similar enhanced susceptibility phenotype and a reduction in *PR1* expression suggests that interference with proteasome function in general leads to defects in immunity. Another report shows that RPT2a is involved in the defence response mediated by a CC-NBS-LRR protein (Chung and Tasaka, 2011). In that case, RPT2a interacts with the resistance protein UNI/uni-1D and a loss of *RPT2a* in the *uni-1D* mutant represses PR1 gene expression, which is normally highly induced in the *uni-1D* mutant plants due to the constitutive activation of the resistance protein (Igari et al., 2008; Chung and Tasaka, 2011).

From a number of additional *Arabidopsis* proteasome subunit mutants tested only those affected in *RPT2a* and *RPN8a* function fully suppressed *edr2*-mediated powdery mildew resistance indicating that the different proteasome subunits might have distinct roles in mediating plant defence responses (Yao et al., 2012). Consistent with this scenario, Hatsugai et al. (2009) reported that the proteasome subunit PBA1 might function as a plant caspase-3-like enzyme, as PBA1 RNAi plants have reduced DEVDase activity besides a decreased overall proteasome activity (Hatsugai et al., 2009). This defect abolished the membrane fusion associated with both disease resistance and HR in response avirulent bacterial strains but not to a virulent strain. As a consequence of this compromised HR, PBA1 RNAi plants display enhanced susceptibility toward avirulent *Pseudomonas* strains, while the growth of virulent *Pst* DC3000 is comparable to *Arabidopsis* wild-type plants (Hatsugai et al., 2009).

A possible explanation for the enhanced susceptibility phenotype of *rpn1a* plants that has been put forward is that a defect in proteasome function interferes with the turnover of a regulator of SA signalling and thus, prevents the onset of SA-mediated defence (Yao et al., 2012). A similar scenario could at least in part explain the compromised immunity in *rpt2a-2* and *rpn12a-1*mutants. A possible candidate for proteasomal turnover during defence is NPR1, the master regulator of SA signalling that acts as a transcriptional co-regulator inside the nucleus and whose functionality was shown to be dependent on continuous proteasomal degradation (Spoel et al., 2009). Compromised NPR1 function due to reduced proteasomal turnover would also explain the reduced expression of the NPR1-downstream target gene *PR1* in proteasome mutants.

However, proteasomal degradation was also reported for immune components acting at early stages of PTI, e.g. the receptor kinase FLS2. Recently, it was revealed that FLS2 is ubiquitinated by the E3 ligases PUB12 and 13 leading to its degradation via the proteasome to attenuate immune responses (Lu et al., 2011). The *pub12* and *pub13* mutants displayed elevated immune responses to flagellin treatment indicating that these E3 ligases act as negative regulators of PTI. The observation that *rpt2a* and *rpn12a* plants show a reduced induction of PTI marker genes *WRKY11* and *WRKY29* suggests that the proteasome can also act as a positive regulator of induced immunity, either at the level of PAMP perception or downstream signalling into the nucleus. Compromised proteasome function could for instance interfere with the turnover of regulators of PTI gene induction. It has been shown in rice (*Oryza sativa*) that WRKY45 degradation via the proteasome is required for the activity of WRKY45 as a transcriptional activator of certain branches of immunity (Matsushita et al., 2013). The likely orthologue of rice WRKY45 is WRKY70 in *Arabidopsis* although proteasomal turnover has so far not been demonstrated to be required for WRKY function in *Arabidopsis* it is conceivable that similar mechanisms are involved in fine tuning defence gene expression also in this species.

The data on the altered kinetics of MAP Kinase phosphorylation further supports the hypothesis that upstream PRR signalling such as FLS2 degradation could be disturbed in the proteasome mutants, but does not exclude direct effects of proteasome activity and ubiquitination on MAPKinase signalling. Generation of ROS also depends on phosphorylation events during the first steps of PAMP recognition. The plasma-membrane-associated kinase BIK1, which is a direct substrate of theFLS2-BAK1 complex, directly interacts with and phosphorylates RBOHD, which is the NADPH oxidase that generates ROS (Kadota et al., 2014). Given the fact that continuous proteasome-mediated degradation of BIK1 ensures optimal immune outputs (Monaghan et al., 2014), it might be possible that the proteasomal turnover of this important immune kinase is affected in the *rpt2a-2* and *rpn12a-1* mutant plants, supporting our findings that ROS production is perturbed in plants with lowered proteasome activity.

Apart from its function in regulating the turnover of components implicated in ROS signalling, proteasome components have been identified to directly contribute to ROS-mediated defence. In tobacco, expression of three genes encoding subunits of the 20S proteasome is induced after treatment with the elicitor cryptogein (Dahan et al., 2001; Suty et al., 2003). Tobacco cell lines overexpressing the ß1 subunit resulted in a decrease of the *NtRbohD* (NADPH oxidase) gene induction and of its associated oxidative burst after cryptogein treatment, indicating that this subunit acts as a negative regulator of early plant responses to cryptogein (Lequeu et al., 2005). Because a loss of *RPT2a* leads to an accumulation of PBA1 (Lee et al., 2011), the homologue of the tobacco ß1 subunit, it is tempting to speculate that ROS signalling is impaired in these plants similar to the overexpressing ß1 tobacco lines.

In addition to a reduced local defence response, Arabidopsis *rpt2a-2* and *rpn12a-1*mutants are also compromised in the establishment of SAR and defence priming towards a secondary infection with bacteria. Mounting of SAR requires the generation of signal in locally infected leaves which is subsequently transmitted to systemic tissue where it is perceived and confers a primed state that enables a faster and stronger defence response upon a secondary infection (Conrath et al., 2015). The exact nature of the signal(s) involved in this process is currently up for debate; however, it is proposed that several hormonal pathways, such as SA, ethylene, auxin and jasmonic acid play a role in SAR (Pastor et al., 2014). Because of its involvement in nearly all aspects of hormonal signalling it appears reasonable to assume that the proteasome could play a critical role for the execution of the different phases of priming. Based on genetic and physiological evidence SA is supposed to play a pivotal role in SAR (Fu and Dong, 2013). SA-signalling strongly depends on NPR1 as a central regulator and interference with the proteasomal turnover of NPR1 would thus also affect SAR and defence priming in *rpt2a-2* and *rpn12a-1* plants. However, recent evidence suggests that the non-protein amino acid pipecolic acid (pip) is a critical SAR regulator (Navarova et al., 2012). Pip elevations are indispensable for SAR and necessary for virtually the whole transcriptional SAR response, although a moderate but significant SA-independent component of SAR activation and SAR gene expression has recently been revealed (Bernsdorff et al., 2016). Future experiments will have to clarify at which step the initiation of SAR is affected by a defect in proteasome function.

Measurement of the overall proteasome activity in leaves infected with a non-pathogenic *Pst* Δ*hrcC*strain revealed a significant induction of proteasome activity. Thus, increased proteasome activity appears to be part of the defence response induced by *Pst*bacteria unable to deliver T3Es into the potential host cell. Consistent with this finding, previous work demonstrated that an *Xcv* strain lacking a functional T3SS also induced proteasome activity upon infection of pepper plants (Üstün et al., 2013). This argues for a contribution of early signalling events, such as recognition of PAMPs and subsequent phosphorylation events, to the induction of the proteasome activity. In accordance with that, the bacterial PAMP flg22 has been shown to activate proteasome peptidase activity upon application leading to alteration in posttranslational modifications in certain proteasome subunits (Sun et al., 2013). Treatment with flg22 also resulted in an accumulation of ubiquitinated proteins (Sun et al., 2013), which is in line with our observation that massive protein turnover leads to an enhanced ubiquitination of proteins during PTI.

The *npr1-1* mutant that is defective in SA-dependent defence responses does not display induction of proteasome activity after infection with non-pathogenic *Pst* suggesting that elevation of proteasomal activity during induced defence depends on the SA-signalling pathway. Previous work implies that SA acts on the transcriptional level to upregulate the expression of certain subunits of the proteasome (Yao et al., 2012; Üstün et al., 2013). In addition, post-translational modification of subunits, e.g. by phosphorylation, would provide a means to rapidly alter the activity or other biochemical properties of the complex.

Virulent *Pst* bacteria that inject the full repertoire of T3Es into their host cell are not only able to prevent induction of proteasome activity, likely through their ability to interfere with SA-mediated defence, but also suppress it below the basal level detected in the mock treated control. This suggests that T3Es act to suppress induction of proteasome activity during defence, which likely occurs at different levels and through different sets of effector proteins. First, the ability to prevent induction of proteasome activity is consistent with the activity of T3Es acting to suppress SA-mediated defence responses (Kazan and Lyons, 2014). In addition, in *npr1-1* wild type *Pst* bacteria not only prevent induction of the proteasome but are also able to further inhibit its activity below the basal threshold independent of interfering with SA-signalling. This indicates that T3Es also exert a more direct effect on the proteasome to reduce its activity. The ability to prevent induction of proteasome activity is independent of the capacity to interfere with very early events of pathogen perception such as the activation of FLS2 by flg22 because a *Pst* mutant lacking the two effectors AvrPto and AvrPtoB is still able to prevent elevated activity levels. Several *Pseudomonas* T3Es that have been implicated to interfere with SA production or signalling. For instance HopI1 has been shown to interfere with SA synthesis inside chloroplasts preventing its accumulation (Jelenska et al., 2007; Jelenska et al., 2010). In addition, HopM1 and AvrE are representatives of T3Es which have the ability to suppress SA-dependent basal immunity and disease necrosis although the targets of these effectors with respect to SA signalling remain to be discovered (DebRoy et al., 2004). However, the interference with SA synthesis *per se* seems not to be sufficient to reduce proteasome activity below basal levels as transient expression of HopI1 in leaves of *N. benthamiana* shows no effect on activity.

A direct inhibition of the proteasome through T3Es of *Pst* targeting its components so far has not been described. However, XopJ a T3E from *X. campestris* pv. *vesicatoria* 85-10 was shown to proteolytically degrade the 19S RP subunit RPT6 in order to inhibit its activity and to interfere with SA-mediated defence in susceptible pepper host plants (Üstün et al., 2013; Üstün and Börnke, 2015). A similar mechanism has been proposed for the T3E HopZ4 from *Pseudomonas syringae* pv. *lachrymans*(Üstün et al., 2014). Both effectors belong to the YopJ-family which is widespread among animal and plant pathogens but whose members are absent from *Pst* (Lewis et al., 2011). Also, *Pst* does not possess SylA, a secreted toxin produced for instance by *P.syringae* pv. *syringae*, which directly targets the catalytic subunits of the 26S proteasome to inhibit its activity and to suppress plant immune reactions, such as SA signalling and stomatal immunity (Baltrus et al., 2011; Groll et al., 2008; Schellenberg et al., 2010; Misas-Villamil et al., 2013). Thus, the suppression of proteasome activity below the basal level by virulent *Pst* likely involves a yet undescribed effector action. We have conducted a systematic screen of a collection of *Pst* T3Es for their ability to suppress proteasome activity when transiently expressed in leaves of *N. benthamiana*. The analyses identified four T3Es, namely HopM1, HopG1, HopAO1 and HopA1 which reproducibly inhibited the proteasome in *N. benthamiana*and thus represent candidates for effectors interfering with proteasome activity during infection of *Arabidopsis* with *Pst*. The function of HopG1, HopAO1 and HopA1 so far has not been shown to be associated with the UPS. HopG1 was demonstrated to inhibit plant innate immunity associated with its localization to mitochondria (Block et al., 2010), while HopA1 associates with EDS1 to disrupt EDS1-TIR NB LRR disease complexes (Bhattacharjee et al., 2011). As the direct target proteins for both T3Es are still not known, it is possible that both T3Es could directly or indirectly associate with the UPS to modulate proteasome activity. Tyrosine phosphatase HopAO1 targets the *Arabidopsis* receptor kinase EF-TU RECEPTOR (EFR) reducing its phosphorylation and thus inhibiting PTI activation (Macho et al., 2014). Because phosphorylation of certain proteasome subunits is crucial to activate its assembly and also induce its activity (Satoh et al., 2001), it might be possible that HopAO1 could target components of the proteasome to reduce their phosphorylation status. Whether this T3E is also able to interact with multiple target proteins in plants, e.g. with components of the proteasome or other UPS-related proteins will be subject of future studies and clarify its role as a proteasome inhibitor.

The identification of HopM1 as a candidate effector protein for suppression of the proteasome is striking. This effect is unlikely to be related to its ability to interfere with SA-dependent defence responses (DebRoy et al., 2004) because a HopM1 deletion strain still prevents induction of proteasome activity above basal levels. However, in contrast to wild type *Pst ΔhopM1* bacteria have lost the ability to suppress proteasome activity below the levels of the mock infected control suggesting that HopM1 is directly interfering with proteasome function. The discovery of HopM1 as a proteasome inhibitor apparently contradicts previous findings, where it has been shown that HopM1 promotes the proteasome-dependent degradation of AtMIN7, a host ADP ribosylation factor guanine nucleotide exchange factor required for vesicle trafficking during PTI and ETI (Nomura et al., 2006; Nomura et al., 2011). However, AP-MS analysis suggests that HopM1 resides in a complex with several proteasome subunits, opening the possibility that HopM1 might directly target proteasome components to interfere with its function. Moreover, using a Y2H approach HopM1 was identified to interact with RAD23, an ubiquitin receptor that delivers ubiquitinated proteins to the 26S proteasome for degradation (Nomura et al., 2006). We speculate that this interaction, on the one hand mediates the degradation of AtMIN7, but on the other hand might affect the recognition of other ubiquitinated proteins by the 26S proteasome. This could result in an inefficient degradation of other substrates and could explain why AtMIN7 is removed by the proteasome while overall proteasome activity is down-regulated. Besides its function to destabilize AtMIN7 and thereby interfering with vesicle trafficking, HopM1 also has an AtMIN7-independent function: it is able to suppress ROS production and stomatal closure during plant immunity (Lozano-Duran et al., 2014). The increased protein levels of FLS2 observed in HopM1 expressing cells supports the hypothesis that impaired proteasome activity may interfere with the proper recycling of the receptor, thereby dampening the PTI response. It is highly plausible that HopM1 triggers the accumulation of inactive FLS2 receptors in the cells by compromising the recycling of the ubiquitinated receptor post elicitation. This effect on basal defence responses is in line with our observations that the proteasome inhibition by genetic means negatively affects ROS production, defence gene expression and also MAPKinase signalling. Thus, these results and previous findings suggest that HopM1 suppresses proteasome activity to dampen ROS generation and suppress stomatal closure to ensure bacterial proliferation during infection.

In conclusion, this work further supports the proposition for a prominent role of the proteasome during early and late defence responses towards Pseudomonas and establishes that *Pst* possess T3Es, which directly or indirectly interfere with proteasome activity during infection. These T3Es may affect proteasome function through multiple mechanisms by (1) preventing the SA-dependent induction of activity above basal levels, and (2) through direct interaction with proteasomal components to interfere with their function. Our experiments have provided candidate T3Es for this second group and future studies will have to clarify the molecular mechanisms of how these effectors target the proteasome.

## Material and Methods

### Plant material and growth conditions

Tobacco plants (*Nicotiana benthamiana*) were grown in soil in a greenhouse with daily watering, and subjected to a 16 h light: 8 h dark cycle (25°C: 21°C) at 300 µmol m–2 s–1 light and 40% relative humidity. *Arabidopsis thaliana* seeds sown on Murashige and Skoog (MS) agar (Sigma-Aldrich) plates supplemented with 2% (w/v) sucrose and cultivated in tissue culture under a 16-h-light/8-h-dark regime (irradiance 150 µmol quanta m^−2^ s^−1^) at 50% humidity. *A. thaliana* seeds germinated on soil were grown under short day conditions (8h light/16h dark [23 °C/ 21 °C]). The *rpt2-2* and *rpn12a-1* mutants in the Col-0 background were originally described in Kurepa et al. 2008.

### Cultivation of Bacteria

*Pseudomonas syringae* pv *maculicola* strain ES4326 (*Psm*) and *Pseudomonas syringae* pv. *tomato* DC3000 (*Pst*) wild-type and deletion strains were grown in King’s B medium containing the appropriate antibiotics at 28°C.

### Assessment of SAR, Defense Priming during SAR, and Local Plant Resistance

The experiments were essentially carried out as detailed in Navarova et al., 2012 with slight modifications. In brief, for SAR induction, plants were infiltrated into three lower (1°) leaves with a suspension of *Psm* at OD 0.005. Infiltration with 10mM MgCl_2_ served as a control treatment. For SAR growth assays, 2° leaves were inoculated with *Psm* (OD 0.001) 2 d after the 1° treatment. Growth of *Psm* in 2° leaves was scored another 2 d later by homogenizing discs originating from infiltrated areas of three different leaves, plating appropriate dilutions on King’s B medium, and counting colony numbers after incubating the plates at 28°C for 2 d.

For the assessment of defence priming during SAR, 2° leaves were infiltrated with either 10 mM MgCl_2_ or *Psm* (OD = 0.005) 2 d after the 1° treatment. The 2° leaves were collected 10 h after the treatment. For the determination of local defence responses such as gene expression of PR1, bacterial suspensions of 1x10^7^ were infiltrated and harvested 24hpi for RNA extraction. For local growth assays bacterial solution of OD 0.002 (*Pst* and *Psm*) were infiltrated into three full-grown leaves per plant. Bacterial growth was assessed 2 d after infiltration as described above.

### Transient expression assays

For infiltration of *N. benthamiana* leaves, *A. tumefaciens* C58C1 was infiltrated into the abaxial air space of 4- to 6-week-old plants, using a needleless 2-ml syringe. Agrobacteria were cultivated overnight at 28°C in the presence of appropriate antibiotics. The cultures were harvested by centrifugation, and the pellet was resuspended in sterile water to a final optical density at (OD600) of 1.0. The cells were used for the infiltration directly after resuspension.

### Western blotting

Leaf material was homogenized in sodium-dodecyl sulphate-polyacrylamide gel electrophoresis (SDS-PAGE) loading buffer (100 mM Tris-HCl, pH 6.8; 9% β-mercaptoethanol, 40% glycerol, 0.0005% bromophenol blue, 4% SDS) and, after heating for 10 min at 95 °C, subjected to gelectrophoresis. Separated proteins were transferred onto nitrocellulose membrane. Proteins were detected by an anti-HA-Peroxidase high affinity antibody (Roche), anti-ubiquitin antibody (Agrisera), anti-AtPBA1 (Enzo life sciences) via chemiluminescence.

### MAP kinase assay

Arabidopsis seedlings were grown for 12 days on MS agar plates and then transferred to 6-well plates (4-6 seedlings per well) in which each well contained 4 ml of liquid medium containing liquid 1xMS media. Seedlings were treated with 1 µ M flg22 peptide and after 0 to 30 min as indicated; the seedlings were frozen in liquid nitrogen. The frozen seedlings were ground in liquid nitrogen and homogenized in 100 µl of extraction buffer (50 mM Tris-HCl, pH 7.5, 100mM NaCl, 15 mM EGTA, 10 mM MgCl_2_, 1mM Na_2_MoO_4_*2H_2_O, 0.5 mM NaVO_3_, 1 mM NaF, 30 mM ß-glycerolphosphate, 0.5 mM PMSF, 1 tablet/10 ml extraction buffer of proteinase inhibitor cocktail (Roche) and phosphatase inhibitor cocktail (Roche)). After centrifugation at 13,000 rpm for 30 min at 4°C, protein concentration of the supernatants was determined using a Bradford assay. Thirty micrograms of protein was separated in an 12,5% polyacrylamide gel. Immunoblot analysis was performed using anti-phospho-p44/42 MAPK (1:1000, Cell Signaling Technology, Danvers, MA, USA) and anti-AtMPK6 (1:2000, Sigma) as primary antibody, and peroxidase-conjugated goat anti-rabbit antibody (1:5,000, Sigma).

### ROS assay

Eight leaf discs (4 mm diameter) from four 4-week-old Arabidopsis plants were sampled using a cork borer and floated overnight on sterile water. The following day the water was replaced with a solution of 17 mg/mL (w/v) luminol (Sigma) and 10 mg/mL horseradish peroxidase (Sigma) containing 1µM flg22. Luminescence was captured over 45min using Synergy HT (BioTek Instruments GmbH) multiplate reader.

### Measurement of proteasome activity

Proteasome activity in plant extracts was determined spectro-fluorometrically using the fluorogenic substrate suc-LLVY-NH-AMC (Sigma) according to Üstün et al 2013.

### RNA Extraction and Expression Analysis

Total RNA was isolated from leafmaterial as described and then treated with RNase-free DNase to degrade any remaining DNA. First-strand cDNA synthesis was performed from 2 µg of total RNA using Revert-Aid reverse transcriptase. For quantitative RT-PCR, the cDNAs were amplified using SensiFAST SYBR Lo-ROX Mix (Bioline GmbH, Luckenwalde) in the AriaMx Realtime PCR System (Agilent Technologies). PCR was optimized, and reactions were performed in triplicate. The transcript level was standardized based on cDNA amplification of *UBC9* (ubiquitin carrier protein) as a reference. Statistical analysis was performed using Student’s t test.

### Protein Extraction and Mass Spectrometry

Adult *Nicotiana benthamiana* plants were syringe infiltrated with Agrobacterium either expressing HopM1-GFP (HopM1 cloned in pGWB5 vector) or a vector GFP alone. After three days, 10g of plant tissue was ground in liquid nitrogen and proteins were isolated in extraction buffer (150 mM Tris, pH 7.5, 150 mM NaCl, 10% glycerol, 5 mM EDTA, 2 mM EGTA, 10 mM DTT, 2% [w/v] polyvinylpyrrolidone, 1% [v/v] Protease Inhibitor Cocktail (PIC, Sigma-Aldrich), 50 mM NaF, 10 mM Na_2_MoO_4_, 1 mM sodium orthovanadate, 0.5 mM phenylmethylsulphonyl fluoride. After centrifugation at 20,000 rcf at 4°C for 30 min, the supernatants were filtered through a miracloth (Millipore). Supernatants were incubated for 2 h at 4°C with 200 ul of anti-GFP trap beads (Chromatek). The beads were washed thrice with 1ml of modified extraction buffer (150 mM Tris, pH 7.5, 150 mM NaCl, 10% glycerol, 5 mM EDTA, 2 mM EGTA and 1% PIC). Proteins were eluted by adding 50 μl of 5x SDS buffer and boiled for 5-10 min and run on 10% SDS-PAGE gel. The gel was stained with Commassie colloidal stain (Invitrogen) and proteins from each lane were trypsin digested and subjected to LC-MS/MS analysis. Reversed phase chromatography was used to separate tryptic peptides prior to mass spectrometric analysis using an Acclaim PepMap µ-precolumn cartridge 300 µm i.d. x 5 mm 5 μm 100 Å and an Acclaim PepMap RSLC 75 µm x 25 cm 2 µm 100 Å (Thermo Scientific). Peptides were eluted directly via a Triversa Nanomate nanospray source (Advion Biosciences, NY) into a Thermo Orbitrap Fusion (Q-OT-qIT, Thermo Scientific) mass spectrometer. The raw data was processed using MSConvert in ProteoWizard Toolkit (version 3.0.5759)1. MS2 spectra were searched with Mascot engine (Matrix Science, version 2.4.1) against *Nicotiana benthamiana* database (supplied by The Sainsbury Laboratory), *Pseudomonas syringae* database (http://www.uniprot.org) and the common Repository of Adventitious Proteins Database (www.thegpm.org/cRAP/index.html). Peptides were generated from a tryptic digestion with up to two missed cleavages, carbamidomethylation of cysteines as fixed modifications, and oxidation of methionines as variable modifications. Precursor mass tolerance was 5 ppm and product ions were searched at 0.8 Da tolerances. Scaffold (version Scaffold_4.3.2, Proteome Software Inc.) was used to validate MS/MS based peptide and protein identifications. Peptide identifications were accepted if they could be established at greater than 95.0% probability by the Scaffold Local FDR algorithm.

## Acknowledgements

We thank Susanne Jeserigk for technical assistance, Alan Collmer and Jürgen Zeier for providing Pseudomonas strains. We are grateful to Jan Smalle for providing the *rpt2a-2* and *rpn12a-1* seeds. We acknowledge the contribution of the Proteomic Research Technology Platform at Warwick University. Vardis Ntoukakis is supported by the Royal Society. This work was partially funded by Biotechnology and Biological Science Research Council grant number BB/L019345/1 to Vardis Ntoukakis.

## References

AlfanoJR, Charkowski AO, Deng WL, Badel JL, Petnicki-Ocwieja T, van DijkK, Collmer A(2000) *The Pseudomonas syringae* Hrp pathogenicity island has a tripartite mosaic structure composed of a cluster of type III secretion genes bounded by exchangeable effector and conserved effector loci that contribute to parasitic fitness and pathogenicity in plants. Proceedings of the National Academy of Sciences of the United States of America 97:4856–4861

Baltrus DA, Nishimura MT, Romanchuk A, Chang JH, Mukhtar MS, Cherkis K, Roach J, Grant SR, Jones CD, Dangl JL (2011) Dynamic evolution of pathogenicity revealed by sequencing and comparative genomics of 19 *Pseudomonas syringae* isolates. PLoS Pathog 7: e1002132

Banfield MJ (2015) Perturbation of host ubiquitin systems by plant pathogen/pest effector proteins. Cell Microbiol 17: 18–25

Bernsdorff F, Doring AC, Gruner K, Schuck S, Brautigam A, Zeier J (2016) Pipecolic Acid Orchestrates Plant Systemic Acquired Resistance and Defense Priming via Salicylic Acid-Dependent and-Independent Pathways. Plant Cell 28: 102–129

Bhattacharjee S, Halane MK, Kim SH, Gassmann W (2011) Pathogen effectors target Arabidopsis EDS1 and alter its interactions with immune regulators. Science 334: 1405–1408

Block A, Guo M, Li G, Elowsky C, Clemente TE, Alfano JR (2010) The *Pseudomonas syringae* type III effector HopG1 targets mitochondria, alters plant development and suppresses plant innate immunity. Cell Microbiol 12: 318–330

Boller T, Felix G (2009) A renaissance of elicitors: perception of microbe-associated molecular patterns and danger signals by pattern-recognition receptors. Annu Rev Plant Biol 60: 379–406

Book AJ, Gladman NP, Lee SS, Scalf M, Smith LM, Vierstra RD (2010) Affinity purification of the Arabidopsis 26 S proteasome reveals a diverse array of plant proteolytic complexes. J Biol Chem 285: 25554–25569

Cao H, Glazebrook J, Clarke JD, Volko S, Dong X (1997) The Arabidopsis NPR1 gene that controls systemic acquired resistance encodes a novel protein containing ankyrin repeats. Cell 88: 57–63

Chung K, Tasaka M (2011) RPT2a, a 26S proteasome AAA-ATPase, is directly involved in Arabidopsis CC-NBS-LRR protein uni-1D-induced signaling pathways. Plant Cell Physiol 52: 1657–1664

Conrath U, Beckers GJ, Langenbach CJ, Jaskiewicz MR (2015) Priming for enhanced defense. Annu Rev Phytopathol 53: 97–119

Cunnac S, Chakravarthy S, Kvitko BH, Russell AB, Martin GB, Collmer A (2011) Genetic disassembly and combinatorial reassembly identify a minimal functional repertoire of type III effectors in *Pseudomonas syringae*. Proc Natl Acad Sci U S A 108: 2975–2980

Dahan J, Etienne P, Petitot AS, Houot V, Blein JP, Suty L (2001) Cryptogein affects expression of alpha3, alpha6 and beta1 20S proteasome subunits encoding genes in tobacco. J Exp Bot 52: 1947–1948

De La Mota-Peynado A, Lee SY, Pierce BM, Wani P, Singh CR, Roelofs J (2013) The proteasome-associated protein Ecm29 inhibits proteasomal ATPase activity and in vivo protein degradation by the proteasome. J Biol Chem 288: 29467–29481

DebRoy S, Thilmony R, Kwack YB, Nomura K, He SY (2004) A family of conserved bacterial effectors inhibits salicylic acid-mediated basal immunity and promotes disease necrosis in plants. Proc Natl Acad Sci U S A 101: 9927–9932

Dudler R (2013) Manipulation of Host Proteosomes as a Virulence Mechanism of Plant Pathogens. Annu Rev Phytopathol

Fu ZQ, Dong X (2013) Systemic acquired resistance: turning local infection into global defense. Annu Rev Plant Biol 64: 839–863

Gomez-Gomez L, Boller T (2000) FLS2: an LRR receptor-like kinase involved in the perception of the bacterial elicitor flagellin in Arabidopsis. Mol Cell 5: 1003–1011

Groll M, Schellenberg B, Bachmann AS, Archer CR, Huber R, Powell TK, Lindow S, Kaiser M, Dudler R (2008) A plant pathogen virulence factor inhibits the eukaryotic proteasome by a novel mechanism. Nature 452: 7551–758

Gruner K, Griebel T, Navarova H, Attaran E, Zeier J (2013) Reprogramming of plants during systemic acquired resistance. Front Plant Sci 4: 252

Gu C, Kolodziejek I, Misas-Villamil J, Shindo T, Colby T, Verdoes M, Richau KH, Schmidt J, Overkleeft HS, van der Hoorn RA (2010) Proteasome activity profiling: a simple, robust and versatile method revealing subunit-selective inhibitors and cytoplasmic, defense-induced proteasome activities. Plant J 62: 160–170

Hatsugai N, Iwasaki S, Tamura K, Kondo M, Fuji K, Ogasawara K, NishimuraM, Hara-Nishimura I (2009) A novel membrane fusion-mediated plant immunity against bacterial pathogens. Genes Dev 23: 2496–2506

He P, Shan L, Lin NC, Martin GB, Kemmerling B, Nurnberger T, Sheen J (2006) Specific bacterial suppressors of MAMP signaling upstream of MAPKKK in Arabidopsis innate immunity. Cell 125: 563–575

Hofius D, Tsitsigiannis DI, Jones JD, Mundy J (2007) Inducible cell death in plant immunity. Semin Cancer Biol 17: 166–187

Igari K, Endo S, Hibara K, Aida M, Sakakibara H, Kawasaki T, Tasaka M (2008) Constitutive activation of a CC-NB-LRR protein alters morphogenesis through the cytokinin pathway in Arabidopsis. Plant J 55: 14–27

Jelenska J, van Hal JA, Greenberg JT (2010) *Pseudomonas syringae* hijacks plant stress chaperone machinery for virulence. Proc Natl Acad Sci U S A 1071: 13177–13182

Jelenska J, Yao N, Vinatzer BA, Wright CM, Brodsky JL, Greenberg JT (2007) A J domain virulence effector of *Pseudomonas syringae* remodels host chloroplasts and suppresses defenses. Curr Biol 17: 499–508

Jones JD, Dangl JL (2006)The plant immune system. Nature 444: 323–329

Kadota Y, Sklenar J, Derbyshire P, Stransfeld L, Asai S, Ntoukakis V, Jones JD, Shirasu K, Menke F, Jones A, Zipfel C (2014) Direct regulation of the NADPH oxidase RBOHD by the PRR-associated kinase BIK1 during plant immunity. Mol Cell 54: 43–55

Kazan K, Lyons R (2014) Intervention of Phytohormone Pathways by Pathogen Effectors. Plant Cell 26: 2285–2309

Kurepa J, Smalle JA (2008) Structure, function and regulation of plant proteasomes. Biochimie 90: 324–335

Kvitko BH, Park DH, Velasquez AC, Wei CF, Russell AB, Martin GB, Schneider DJ, Collmer A (2009) Deletions in the repertoire of *Pseudomonas syringae* pv. *tomato* DC3000 type III secretion effector genes reveal functional overlap among effectors. PLoS Pathog 5: e1000388

Lee KH, Minami A, Marshall RS, Book AJ, Farmer LM, Walker JM, Vierstra RD (year>2011) The RPT2 subunit of the 26S proteasome directs complex assembly, histone dynamics, and gametophyte and sporophyte development in Arabidopsis. Plant Cell 23: 4298–4317

Lequeu J, Simon-Plas F, Fromentin J, Etienne P, Petitot AS, Blein JP, Suty L (2005) Proteasome comprising a beta1 inducible subunit acts as a negative regulator of NADPH oxidase during elicitation of plant defense reactions. FEBS Lett 579: 4879–48861

Lewis JD, Lee A, Ma W, Zhou H, Guttman DS, Desveaux D (2011) The YopJ superfamily in plant-associated bacteria. Mol Plant Pathol 12: 928–937

Lozano-Duran R, Bourdais G, He SY, Robatzek S (2014) The bacterial effector HopM1 suppresses PAMP-triggered oxidative burst and stomatal immunity. New Phytol 202: 259–269

Lu D, Lin W, Gao X, Wu S, Cheng C, Avila J, Heese A, Devarenne TP, He P, Shan L (2011) Direct ubiquitination of pattern recognition receptor FLS2 attenuates plant innate immunity. Science 332: 1439–1442

Macho AP (2016) Subversion of plant cellular functions by bacterial type-III effectors: beyond suppression of immunity. New Phytol 210: 51–57

Macho AP, Schwessinger B, Ntoukakis V, Brutus A, Segonzac C, Roy S, Kadota Y, Oh MH, Sklenar J, Derbyshire P, Lozano-Duran R, Malinovsky FG, Monaghan J, Menke FL, Huber SC, He SY, Zipfel C (2014) A bacterial tyrosine phosphatase inhibits plant pattern recognition receptor activation. Science 343: 1509–1512

Marino D, Peeters N, Rivas S (2012) Ubiquitination during plant immune signaling. Plant Physiol 160: 15–27

Matsushita A, Inoue H, Goto S, Nakayama A, Sugano S, Hayashi N, TakatsujiH (2012) The nuclear ubiquitin proteasome degradation affects WRKY45 function in the rice defense program. Plant J

Misas-Villamil JC, Kolodziejek I, Crabill E, Kaschani F, Niessen S, Shindo T, Kaiser M, Alfano JR, van der Hoorn RA (2013) *Pseudomonas syringae* pv. *syringae* uses proteasome inhibitor syringolin A to colonize from wound infection sites. PLoS Pathog 9: e1003281

Mishina TE, Zeier J (2006) The Arabidopsis flavin-dependent monooxygenase FMO1 is an essential component of biologically induced systemic acquired resistance. Plant Physiol 141: 1666–16751

Monaghan J, Matschi S, Shorinola O, Rovenich H, Matei A, Segonzac C, Malinovsky FG, Rathjen JP, MacLean D, Romeis T, Zipfel C (2014) The calcium-dependent protein kinase CPK28 buffers plant immunity and regulates BIK1 turnover. Cell Host Microbe 16: 605–615

Navarova H, Bernsdorff F, Doring AC, Zeier J (2012) Pipecolic acid, an endogenous mediator of defense amplification and priming, is a critical regulator of inducible plant immunity. Plant Cell 24: 5123–5141

Nicaise V, Roux M, Zipfel C (2009) Recent advances in PAMP-triggered immunity against bacteria: pattern recognition receptors watch over and raise the alarm. Plant Physiol 150: 1638–1647

Nomura K, Debroy S, Lee YH, Pumplin N, Jones J, He SY (2006) A bacterial virulence protein suppresses host innate immunity to cause plant disease. Science 313: 220–223

Nomura K, Mecey C, Lee YN, Imboden LA, Chang JH, He SY (2011) Effector-triggered immunity blocks pathogen degradation of an immunity-associated vesicle traffic regulator in Arabidopsis. Proc Natl Acad Sci U S A 108: 10774–10779

Pastor V, Balmer A, Gamir J, Flors V, Mauch-Mani B (2014) Preparing to fight back: generation and storage of priming compounds. Front Plant Sci 5: 295

Sadanandom A, Bailey M, Ewan R, Lee J, Nelis S (2012) The ubiquitin-proteasome system: central modifier of plant signalling. New Phytol 196: 13–28

Satoh K, Sasajima H, Nyoumura KI, Yokosawa H, Sawada H (2001) Assembly of the 26S proteasome is regulated by phosphorylation of the p45/Rpt6 ATPase subunit. Biochemistry 40: 314–319

Schellenberg B, Ramel C, Dudler R (2010) *Pseudomonas syringae* virulence factor syringolin A counteracts stomatal immunity by proteasome inhibition. Mol Plant Microbe Interact 23: 1287–1293

Singer AU, Schulze S, Skarina T, Xu X, Cui H, Eschen-Lippold L, Egler M, Srikumar T, Raught B, Lee J, Scheel D, Savchenko A, Bonas U (2013) A pathogen type III effector with a novel E3 ubiquitin ligase architecture. PLoS Pathog 9: e1003121

Smalle J (2002) Cytokinin Growth Responses in Arabidopsis Involve the 26S Proteasome Subunit RPN12. The Plant Cell Online 14: 17–32

Smalle J, Vierstra RD (2004) The ubiquitin 26S proteasome proteolytic pathway. Annu Rev Plant Biol 55: 555–590

Spoel SH, Mou Z, Tada Y, Spivey NW, Genschik P, Dong X (2009) Proteasome-mediated turnover of the transcription coactivator NPR1 plays dual roles in regulating plant immunity. Cell 137: 860–872

Stegmann M, Anderson RG, Ichimura K, Pecenkova T, Reuter P, Zarsky V, McDowell JM, Shirasu K, Trujillo M (2012) The ubiquitin ligase PUB22 targets a subunit of the exocyst complex required for PAMP-triggered responses in Arabidopsis. Plant Cell 24: 4703–4716

Sun HH, Fukao Y, Ishida S, Yamamoto H, Maekawa S, Fujiwara M, Sato T, Yamaguchi J (2013) Proteomics analysis reveals a highly heterogeneous proteasome composition and the post-translational regulation of peptidase activity under pathogen signaling in plants. J Proteome Res 12: 5084–5095

Suty L, Lequeu J, Lançon A, Etienne P, Petitot A-S, Blein J-P (2003) Preferential induction of 20S proteasome subunits during elicitation of plant defense reactions: towards the characterization of "plant defense proteasomes". The International Journal of Biochemistry & Cell Biology 35: 637–650

Trujillo M, Ichimura K, Casais C, Shirasu K (2008) Negative regulation of PAMP-triggered immunity by an E3 ubiquitin ligase triplet in Arabidopsis. Curr Biol 18: 1396–1401

Üstün S, Bartetzko V, Börnke F (2013) The *Xanthomonas campestris* Type III Effector XopJ Targets the Host Cell Proteasome to Suppress Salicylic-Acid Mediated Plant Defence. PLoS Pathog 9: e1003427

Üstün S, Börnke F (2014) Interactions of Xanthomonas type-III effector proteins with the plant ubiquitin and ubiquitin-like pathways. Front Plant Sci 5: 736

Üstün S, Börnke F (2015) The *Xanthomonas campestris* type III effector XopJ proteolytically degrades proteasome subunit RPT6. Plant Physiol 168: 107–119

Üstün S, König P, Guttman DS, Börnke F (2014) HopZ4 from *Pseudomonas syringae*, a Member of the HopZ Type III Effector Family from the YopJ Superfamily, Inhibits the Proteasome in Plants. Mol Plant Microbe Interact 27: 611–623

Vierstra RD (2009) The ubiquitin-26S proteasome system at the nexus of plant biology. Nat Rev Mol Cell Biol 10: 385–397

Yao C, Wu Y, Nie H, Tang D (2012) RPN1a, a 26S proteasome subunit, is required for innate immunity in Arabidopsis. Plant J 71: 1015–1028

