## Supplemental Figures for "The proteasome acts as a hub for local and systemic plant immunity in *Arabidopsis thaliana* and constitutes a virulence target of *Pseudomonas syringae* type-III effector proteins"

**Supplementary Figure 1**

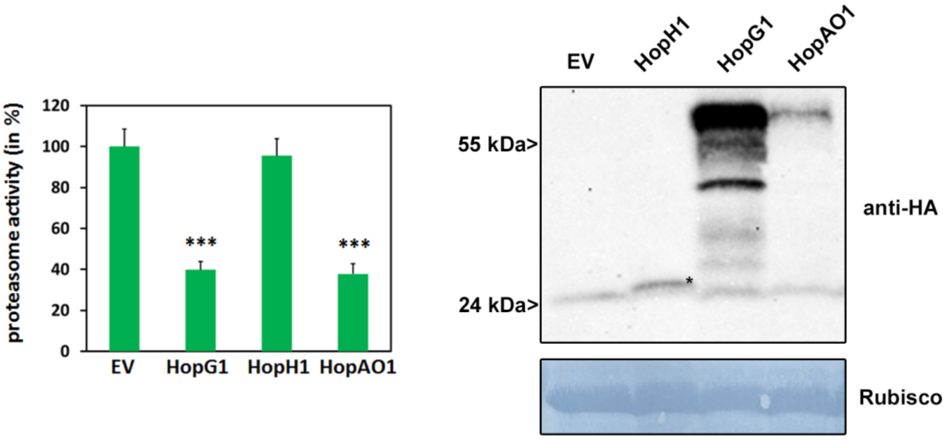


**Figure S1:** Proteasome activity in *N. benthamiana* leaves following transient expression of T3Es HopG1, HopH1, HopAO1 or empty vector control (EV). Relative proteasome activity in total protein extracts was determined by monitoring the breakdown of the fluorogenic peptide suc-LLVY-AMC at 30°C in a fluorescence spectrophotometer. EV was set to 100%. Data represent the mean ± standard deviation (SD) (n = 3). Asterisks indicate statistical significance (***P<0.001) determined by Student´s t test (compared with EV control). Protein extracts from *N. benthamiana* leaves transiently expressing T3Es tagged with HA and empty vector (EV) at 48 hpi were prepared. Equal volumes representing approximately equal protein amounts of each extract were immunoblotted and proteins were detected using anti-HA antiserum. Amido black staining served as a loading control.

**Supplementary Figure 2**

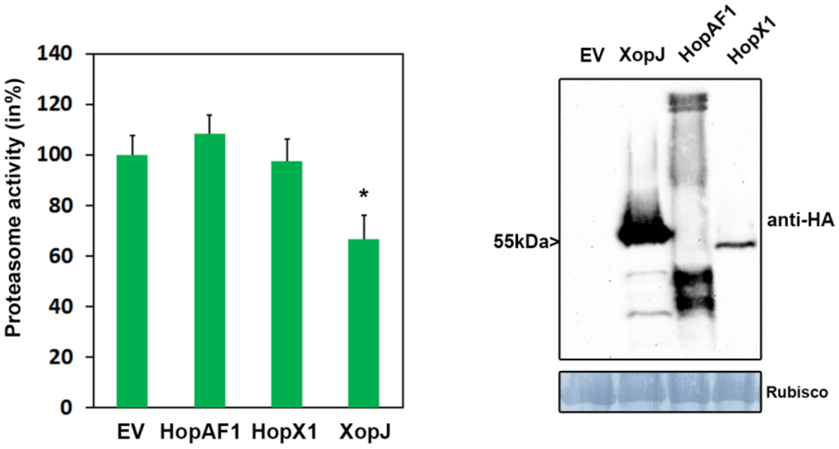


**Figure S2:** Proteasome activity in *N. benthamiana* leaves following transient expression of T3Es HopAF1, HopX1, XopJ or empty vector control (EV). Relative proteasome activity in total protein extracts was determined by monitoring the breakdown of the fluorogenic peptide suc-LLVY-AMC at 30°C in a fluorescence spectrophotometer. EV was set to 100%. Data represent the mean ± standard deviation (SD) (n = 3). Asterisks indicate statistical significance (*P<0.05) determined by Student´s t test (compared with EV control). Protein extracts from *N. benthamiana* leaves transiently expressing T3Es tagged with HA and empty vector (EV) at 48 hpi were prepared. Equal volumes representing approximately equal protein amounts of each extract were immunoblotted and proteins were detected using anti-HA antiserum. Amido black staining served as a loading control.

**Supplementary Figure 3**

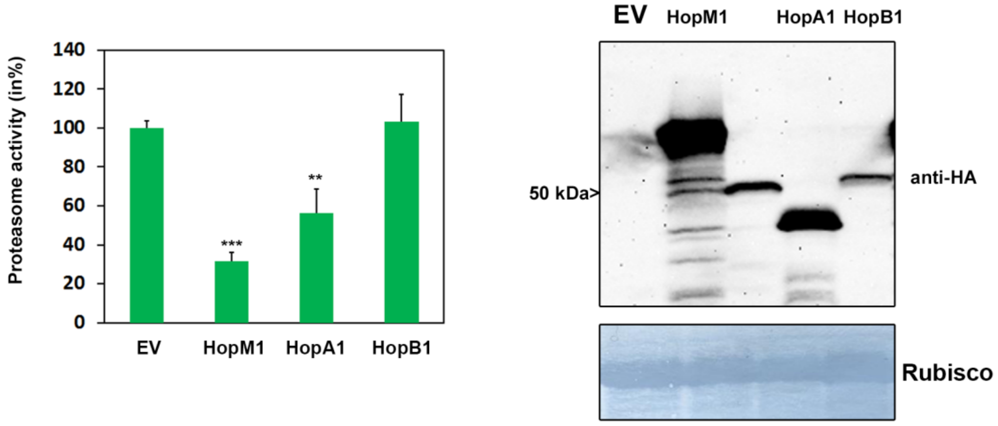


**Figure S3:** Proteasome activity in *N. benthamiana* leaves following transient expression of T3Es HopM1, HopA1, HopB1 or empty vector control (EV). Relative proteasome activity in total protein extracts was determined by monitoring the breakdown of the fluorogenic peptide suc-LLVY-AMC at 30°C in a fluorescence spectrophotometer. EV was set to 100%. Data represent the mean ± standard deviation (SD) (n = 3). Asterisks indicate statistical significance (**P<0.01; ***P<0.001) determined by Student´s t test (compared with EV control). Protein extracts from *N. benthamiana* leaves transiently expressing T3Es tagged with HA and empty vector (EV) at 48 hpi were prepared. Equal volumes representing approximately equal protein amounts of each extract were immunoblotted and proteins were detected using anti-HA antiserum. Amido black staining served as a loading control.

**Supplementary Figure 4**

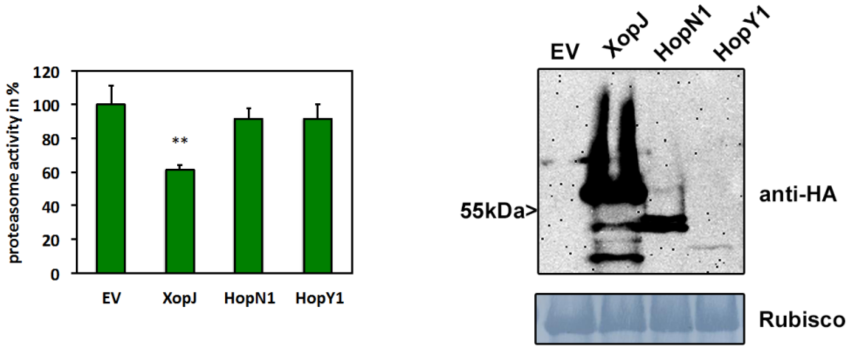


**Figure S4:** Proteasome activity in *N. benthamiana* leaves following transient expression of T3Es HopN1, HopY1, XopJ or empty vector control (EV). Relative proteasome activity in total protein extracts was determined by monitoring the breakdown of the fluorogenic peptide suc-LLVY-AMC at 30°C in a fluorescence spectrophotometer. EV was set to 100%. Data represent the mean ± standard deviation (SD) (n = 3). Asterisks indicate statistical significance (**P<0.01) determined by Student´s t test (compared with EV control). Protein extracts from *N. benthamiana* leaves transiently expressing T3Es tagged with HA and empty vector (EV) at 48 hpi were prepared. Equal volumes representing approximately equal protein amounts of each extract were immunoblotted and proteins were detected using anti-HA antiserum. Amido black staining served as a loading control.

**Supplementary Figure 5**

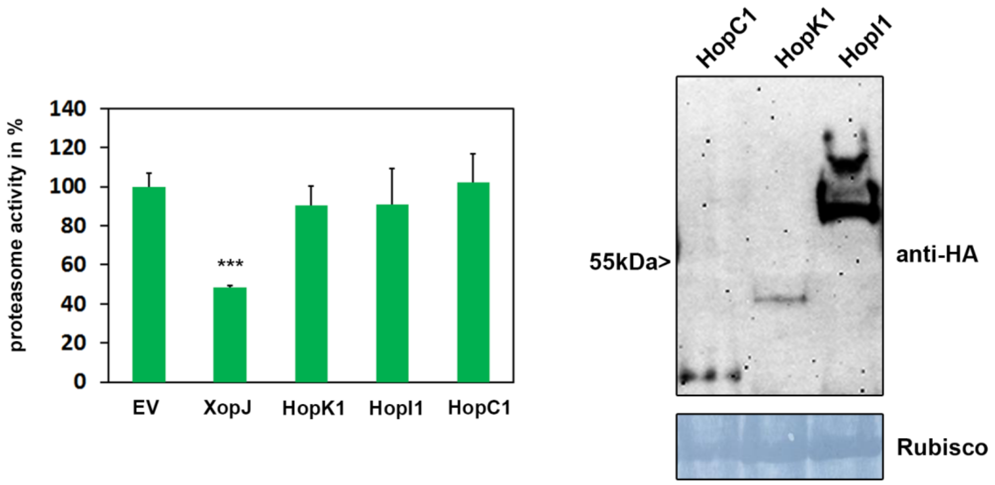


**Figure S5:** Proteasome activity in *N. benthamiana* leaves following transient expression of T3Es HopK1, HopI1, HopC1, XopJ or empty vector control (EV). Relative proteasome activity in total protein extracts was determined by monitoring the breakdown of the fluorogenic peptide suc-LLVY-AMC at 30°C in a fluorescence spectrophotometer. EV was set to 100%. Data represent the mean ± standard deviation (SD) (n = 3). Asterisks indicate statistical significance (***P<0.001) determined by Student´s t test (compared with EV control). Protein extracts from *N. benthamiana* leaves transiently expressing T3Es tagged with HA and empty vector (EV) at 48 hpi were prepared. Equal volumes representing approximately equal protein amounts of each extract were immunoblotted and proteins were detected using anti-HA antiserum. Amido black staining served as a loading control.

**Supplementary Figure 6**

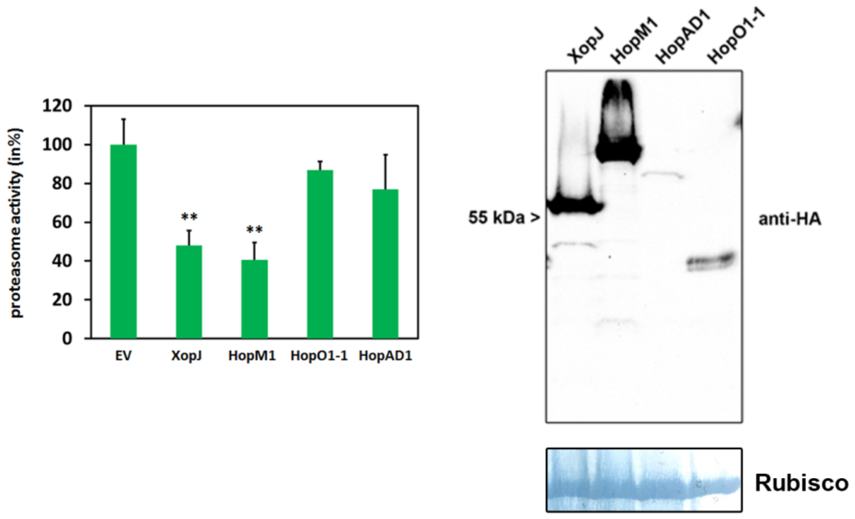


**Figure S6**: Proteasome activity in *N. benthamiana* leaves following transient expression of T3Es HopM1, HopO1-1, HopAD1, XopJ or empty vector control (EV). Relative proteasome activity in total protein extracts was determined by monitoring the breakdown of the fluorogenic peptide suc-LLVY-AMC at 30°C in a fluorescence spectrophotometer. EV was set to 100%. Data represent the mean ± standard deviation (SD) (n = 3). Asterisks indicate statistical significance (**P<0.01) determined by Student´s t test (compared with EV control). Protein extracts from *N. benthamiana* leaves transiently expressing T3Es tagged with HA and empty vector (EV) at 48 hpi were prepared. Equal volumes representing approximately equal protein amounts of each extract were immunoblotted and proteins were detected using anti-HA antiserum. Amido black staining served as a loading control.

**Supplementary Figure 7**

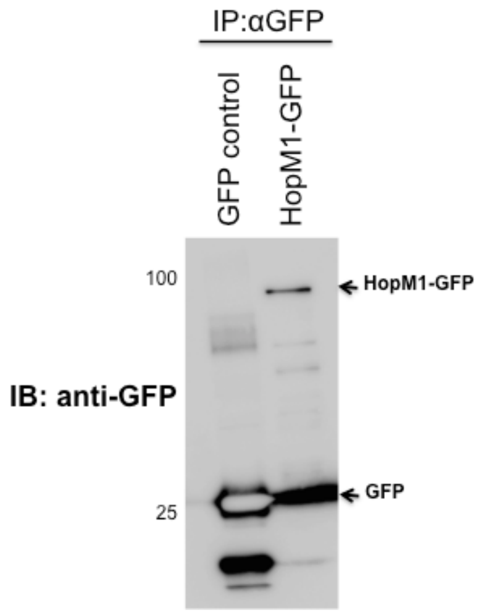


**Figure S7:** HopM1 interacts with proteasome-associated proteins. Immunoprecipitation of HopM1 using GFP trap agarose beads. Western blot using anti-GFP antibody showing the GFP control and C-terminal GFP tagged HopM1 pulled down proteins expressed in *Nicotiana benthamiana*. The arrowheads indicate the position of the proteins. The gel lanes were excised and Mass Spec analysis was performed.
